# Seed-Guided *De Novo* Design Expands the Structural Diversity of Antitoxin Protein Binders

**DOI:** 10.64898/2026.08.02.742339

**Authors:** Dustin Britton, Dia A. Ghose, Jackson C. Halpin, Foster Birnbaum, Kruthi Gundu, Shaunak Raval, Malvina Papanastasiou, Steven A. Carr, Amy E. Keating

## Abstract

*De novo* design of protein binders targeting extended, multi-site interaction surfaces remains difficult for current generative methods, which often produce limited structural diversity and predominantly helical topologies. Here, we advance diffusion-based binder design by guiding protein backbone generation through “seeds,” which are PDB-derived fragments selected for geometric complementarity to the target surface. To test this approach, we computationally generated seed-guided binders of the bacterial toxin RelE. RelB, the native antitoxin of RelE, engages two distinct interfaces with high surface complementarity, making it an appropriate test case. Seed-guided RFdiffusion produced backbones with substantially higher structural diversity and more target contacts than RFdiffusion alone. Experimental screening of 1,402 designs in a high-throughput bacterial survival assay identified multiple functional binders, including variants with nanomolar to low-micromolar affinity and one design with RelE neutralization comparable to RelB_pep_. Computational structure prediction and mutational analyses support that the designed interfaces rely on seed-derived contacts and adopt binding modes distinct from RelB. Molecular dynamics simulations and hydrogen-deuterium exchange experiments further suggest that one high-affinity design undergoes a conformational change upon binding. Notably, successful designs exhibited reduced cross-reactivity to RelE orthologs compared with RelB_pep_, suggesting that the extensive interfaces generated through seed-guided design can enable enhanced selectivity. These results establish motif scaffolding of surface-complementing seeds as an effective strategy for overcoming current limitations of *de novo* generative models, enabling the design of proteins that can engage challenging interface sites.

## Introduction

Nature has evolved a vast repertoire of structurally diverse protein-protein interfaces to scaffold complexes, transduce information, and stimulate or inhibit enzymatic activity (1). Computational design methods, which have advanced rapidly due to deep learning innovations, can now generate *de novo* protein binders for many natural targets, opening new possibilities in biomedicine (2, 3). Interestingly, the binders generated by state-of-the-art generative deep learning models, including diffusion and flow-matching methods, do not closely resemble evolved protein structures. The structural diversity and binding modes of *de novo* designs remain relatively limited (4, 5). Specifically, designed binders rely more heavily on regular secondary structure elements, particularly alpha helices, to stabilize their folds and interactions than natural binders do. Another feature of *de novo* protein binders is that they often make contacts across a relatively limited area of the target protein surface, using a few strong “hot spots” to achieve binding. The structural biases of designed binders may limit the range of applications and functions that can be engineered, including the ability to bind a target through multiple sites. Multi-site binding is common in biology: antibodies use multiple complementarity-determining regions to engage different sites on antigens (6), enzyme inhibitors often occupy extended active sites and adjacent regulatory surfaces (7, 8), and scaffold proteins simultaneously contact multiple partners across distinct interfaces (9–11).

We sought to design *de novo* binders that engage a target across an extensive interface with high structural and chemical complementarity. Our approach leverages structural “seeds,” which are mined from known protein structures in the PDB based on their geometric complementarity to a target surface of interest. Seeds are short peptide backbone fragments derived from tertiary motifs (TERMs) that define recurring spatial arrangements of residues in three-dimensional space, independent of their sequence context (12). Because seeds are derived from naturally occurring residue arrangements with favorable local packing, they provide a physically grounded starting geometry for interface design. Swanson et al. previously developed methods to search structures in the Protein Data Bank (PDB) and identify seeds that complement any target protein. We postulated that these seeds could anchor full-length proteins generated by RFdiffusion, thereby embedding validated binding surfaces into folded protein structures.

As our target, we chose the bacterial toxin RelE, a ribosome-dependent endoribonuclease that is part of a type II toxin-antitoxin (TA) system (**Figure 1a**) (13, 14). TA systems serve as bacterial defense mechanisms against phage infection through abortive infection pathways (15), making selective toxin inhibition potentially valuable for improving bacteriophage-based therapies (16–18). RelE presents an exceptional structural challenge for protein binder design. The native antitoxin RelB neutralizes RelE through a dual binding mode: its N-terminal α-helix targets one interface while its C-terminal β-strand engages a second, spatially distinct interface. Together, these elements wrap around the toxin surface to create extensive contacts and disrupt the enzyme active site (**Figure 1b**). A peptide derived from RelB, here called RelB_pep_ (residues K47–L79), functions as an inhibitor, but the multi-site engagement is essential. Neither the N-terminal nor the C-terminal regions of RelB_pep_ alone inhibit RelE or rescue bacterial growth (19).

**Figure 1.**
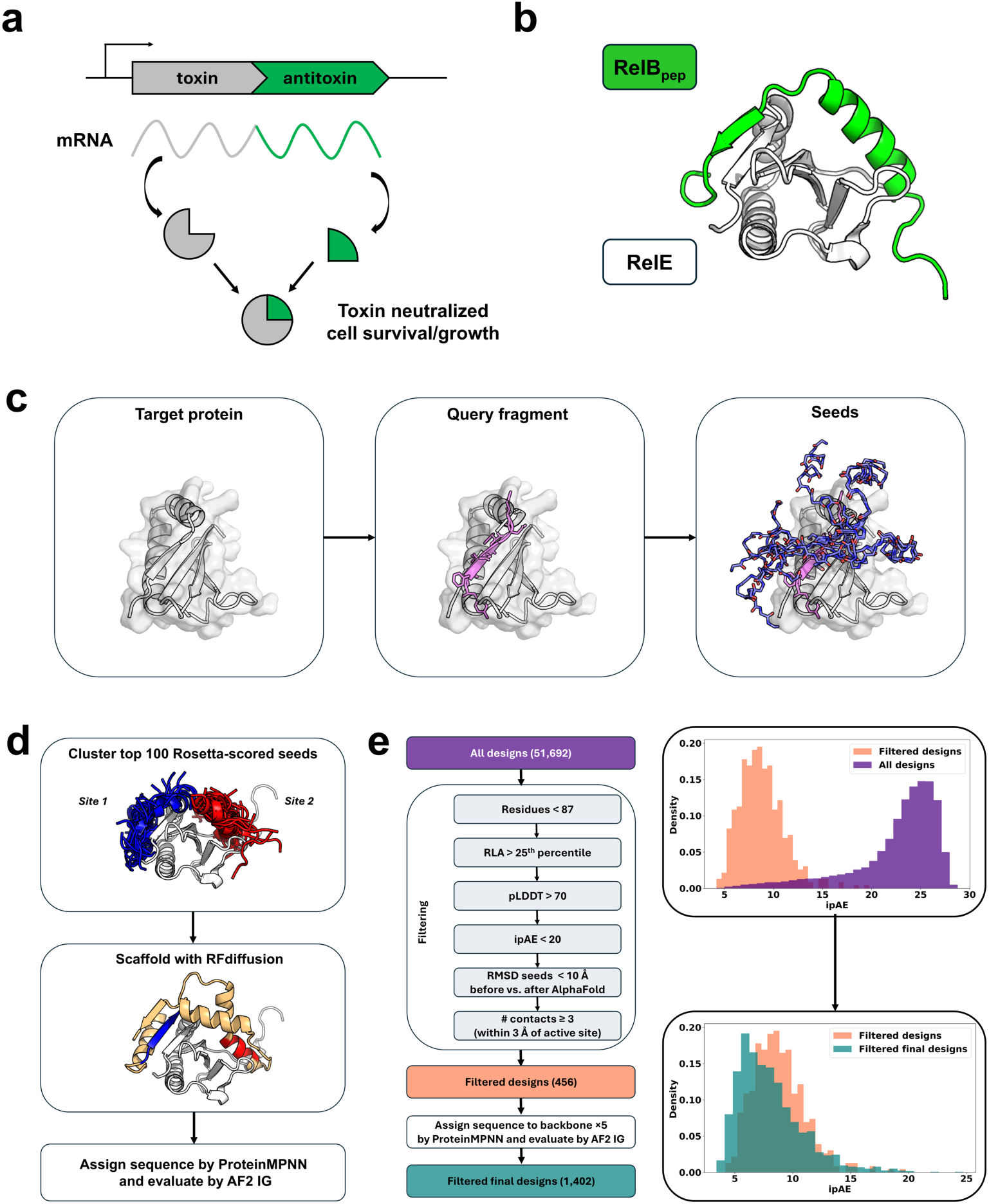
Overview of the computational design of binders of the *E. coli* toxin protein RelE. (**a**) Scheme illustrating how co-operonic toxin-antitoxin pairs impact bacterial cell survival and growth. **(b)** *E. coli* RelE in complex with a portion of RelB, RelB_pep (_PDB: 2KC8) (22). (**c**) A search of the PDB using query fragments from RelE (one example is shown in pink) generated backbone seeds for binder design (shown in blue sticks), as described in the Methods. (**d**) The 100 top-scoring seeds were divided into two clusters based on their RelE binding sites and incorporated into longer protein backbones using RFdiffusion prior to sequence design using ProteinMPNN. (**e**) Pipeline for filtering designs based on AF2 IG structures. ipAE score distributions for designs at different stages of the design and filtering process.

Ghose et al. previously tested multiple strategies for designing RelE binders based on RelE-complementing seeds (19). In one approach, they computationally fused multiple seeds, but none of ∼5,000 linear peptides designed in this way showed enrichment in functional screens. A more successful strategy involved computationally designing a peptide that fused two seeds to the native RelB N-terminal α-helix. This approach yielded 75 peptides with RelE neutralization activity, corresponding to 7.5% of the tested designs of this type. These results suggested that seeds can provide productive binding geometries but lack adequate scaffolding to generate functional proteins.

Here, we return to seed-guided design with an improved strategy: using the motif-scaffolding protocol of RFdiffusion to construct protein backbones around PDB-derived seeds. This approach combines the structural diversity accessible through tertiary motif searching with the powerful backbone generation capabilities of diffusion models. We generated ∼1,400 binders with substantially greater structural diversity and more extensive target contacts than previous approaches. Functional screening in *Escherichia coli* and biophysical characterization revealed multiple hits with neutralization activity and strong binding affinity, validating seed-guided design as an effective strategy for targeting challenging protein-protein interfaces.

## Results

### Seed-Guided *De Novo* Design

To establish a baseline, we first designed binders without seeds. We designed 12,000 protein binders of RelE using RFdiffusion guided by hotspots but not seeds, followed by sequence design with ProteinMPNN (see Methods for details). When evaluated using the AlphaFold2 Initial Guess (AF2 IG) protocol (20, 21), which is an established filter for *de novo* binder design, many designs had high predicted local distance difference test (pLDDT) scores and low interface predicted aligned error (ipAE) scores, indicating that the designed sequences were confidently predicted to adopt the intended fold and interface geometry (**Figure 2a**, **Figure S1**). However, the hotspot-only designs did not make extensive contacts with the target domain. All designs with ipAE < 10 were predicted to locally engage the β-strand hotspot of RelE (involving RelE residues L5, D6, F7, and D8). This region, which we refer to as site 1, is bound by the C-terminal β-strand of RelBpep. However, only 146 (1.2%) designs had confidently predicted interfaces (ipAE < 10) and met or exceeded the number of contacts made between RelB and RelE (PDB: 2KC8; 26 contacts with atoms within 4 Å) (22). Furthermore, only 10 (< 0.1%) of the 146 designs also had at least one contact to the active site, which is a feature of the binding mode of RelB (22) (**Figure 1b**) and a predicted feature of computationally designed RelB helix-extension peptides that inhibit RelE function (19). We also applied BindCraft (23), using the hotspots option, to design RelE binders. Despite the high computational cost (∼ 8.1 days of NVIDIA L40S GPU time), only 502 binders based on 291 backbones passed the default filters. Of these, 29 binders showed an ipAE < 10 and met or exceeded the number of contacts between RelB and RelE (**Figure S2**). None of these designs contacted active-site residues (measured by all-atom distance within 3 Å of residues R45, K52, and R61 in RelE (22)).

**Figure 2.**
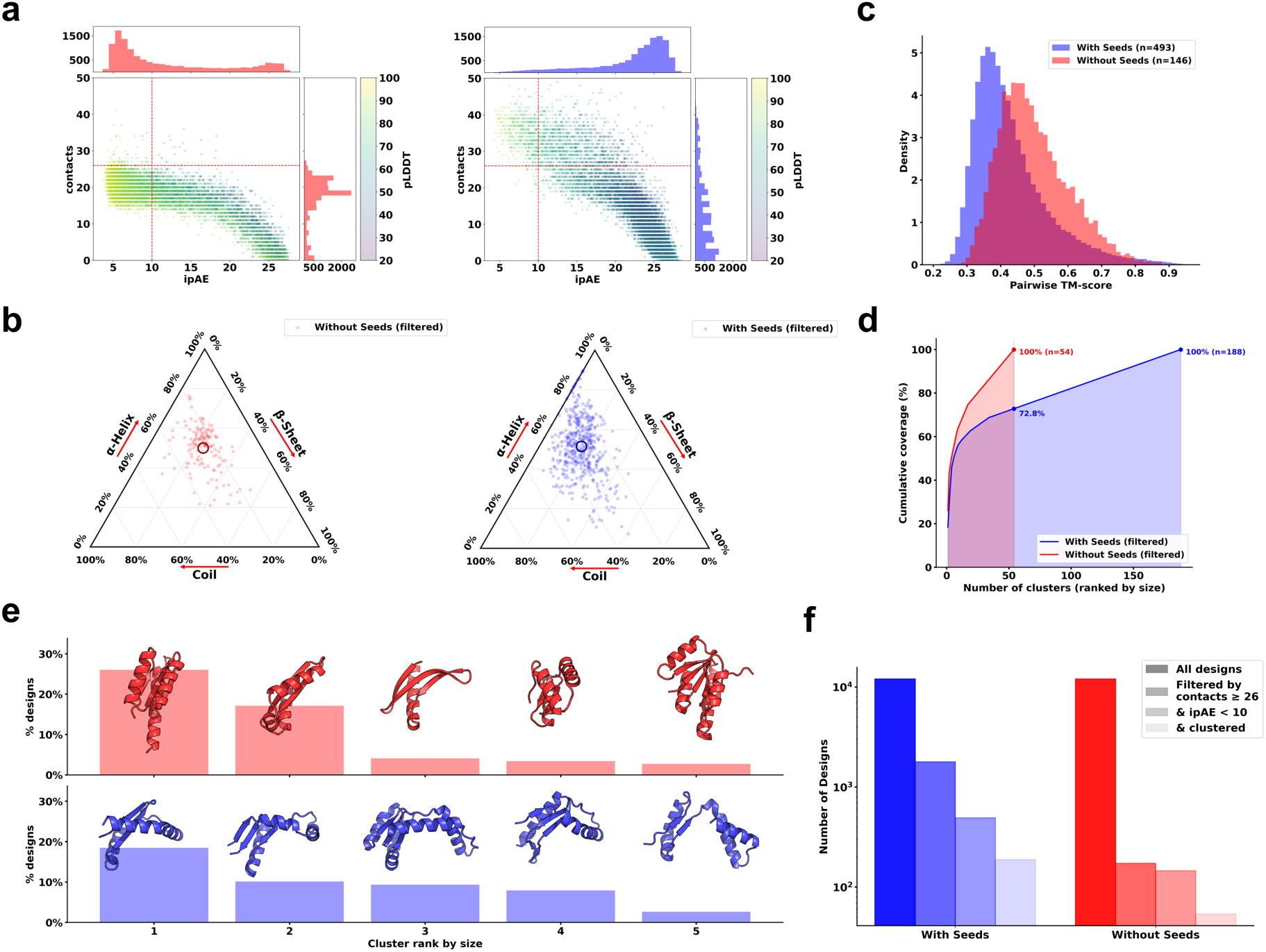
RFdiffusion designs generated using seeds show increased structural diversity compared to those without seeds. We analyzed the structures for 12,000 designs generated without seeds and a corresponding subset of 12,000 structures designed using seeds. (**a**) Designs using seeds had higher ipAE and lower pLDDT confidence scores but made more contacts with RelE (distance < 4 Å). (**b**) Designs using seeds had a greater range of secondary structure compositions calculated with STRIDE (24). The mean secondary structure of all designs is represented with open circles. (**c**) Filtered seed-guided designs (n = 493, blue) had a lower mean pairwise TM-score compared with filtered default RFdiffusion designs (n = 146, red), among designs with ipAE < 10 and contacts ≥ 26. (**d**) Seed-guided designs produced more clusters than no-seed designs and showed less cumulative coverage at an equal number of clusters (73% at 54 clusters for seed-guided designs). (**e**) Clustering with a TM-align threshold of 0.5 gave larger clusters for designs generated without seeds than with seeds, reflecting less diversity. Representative structures are shown for the five largest clusters for designs generated with and without seeds after aligning RelE (not shown), matching the orientation shown in Figure 1b. (**f**) The seed-guided design protocol generated more unique structures than the no-seed protocol, following multiple steps of filtering for high interface confidence and ≥ 26 RelE contacts.

To generate designs with more extensive surface complementarity to RelE, we used seeds to define interfaces and scaffolded them with RFdiffusion. Ghose et al. placed 10,229 seeds around the surface of RelE and scored them by a sequence-structure compatibility score (**Supplementary Materials and Methods**, **Figure 1c)**. The 100 top-scoring seeds from that work engaged RelE in diverse ways but clustered around two parts of the surface that we define as site 1 (53 seeds) and site 2 (47 seeds) (**Figure 1d** and **Figure S3**). Both sites engage native RelB, although knowledge of the binding mode of RelB was not used in seed generation: site 1 corresponds to the interaction site formed by the C-terminal β-strand of RelB_pep_, and site 2 corresponds to the interaction site formed by the N-terminal helix of RelB_pep_. From these clusters, we generated 4,982 ordered pairs by selecting one seed from each site, in both possible orderings, and used RFdiffusion with base or beta weights and the motif-scaffolding protocol to generate 5–6 backbones per pair that scaffolded the fragments in the indicated order (**Figure 1d**). Generating one sequence per backbone with ProteinMPNN gave 51,692 proteins for which we evaluated the structure of the designed binder–RelE complex using AF2 IG (**Figure 1e**) (20).

**Figure 2** compares the unfiltered seed-guided and no-seed designs. For this analysis, we randomly sampled 12,000 seed-guided designs generated with the same RFdiffusion model weights and length distribution as the no-seed designs. Both methods generated designs with a wide range of ipAE scores from < 5 to > 25, though the median confidence of the seed-guided designs was lower (with higher ipAE scores) than that of the no-seed designs (median ipAE = 23.8 vs. 8.5; **Figures 2a and 2b**). Notably, the highest-confidence seed-guided designs, those with ipAE < 10 (examples are shown in **Figure S4**), had very different interface characteristics than the no-seed designs (**Figures S1, S2**). The high-confidence seed-guided designs made many more contacts with RelE (median contacts 35 versus 20, **Figure 2a**). Using combined quality thresholds of ipAE < 10, pLDDT > 70, and ≥ 26 contacts, 4.1% of seed-guided designs passed all three criteria, compared with only 1.2% of no-seed designs. Mapping the frequency of contacts with RelE showed greater coverage across the surface complemented by native RelB for seed-guided designs than for no-seed designs (**Figure S5**). These features of the designs are consistent with the seed-guided approach generating binders that wrap around RelE to engage both the site 1 and site 2 interfaces.

To focus our analysis on the most promising structures, we selected the designs that were confidently predicted to make extensive contacts with RelE. After filtering for those designs with ipAE < 10 that, like RelB, made ≥ 26 contacts with RelE, seed-guided RFdiffusion produced more designs than did no-seed RFdiffusion (493 vs. 146) (**Figure 2a**). BindCraft produced only 29 designs that passed these filters (**Figure S2a**). **Figures 2b and S2b** show the secondary structure compositions of these designs and indicate the average secondary structure for these sets. The distributions of the structures in these plots, and the mean secondary structure content for all designs, highlight the higher coil content in the seed-guided designs than in the no-seed RFdiffusion and BindCraft designs (mean values of 30.1% for seed-guided, 25.7% for no-seed RFdiffusion, and 20.8% for BindCraft).

We calculated pairwise TM-scores for designs in the seed-guided and no-seed sets, which showed a lower average structural similarity (lower TM-score) for the seed-guided designs (**Figure 2c**). Clustering with a TM-align score cutoff of 0.5 produced 188 clusters for the seed-guided designs compared with only 54 clusters for the default RFdiffusion set, also consistent with the seed-guided protocol generating greater diversity (**Figure 2c**). **Figure 2e** shows representative structures for the five largest clusters. For default RFdiffusion, these five clusters contained 3–26% of the high-confidence, high-contact-number structures per cluster, with all five clusters corresponding to a compact, mostly helical architecture with a β-strand contacting the β-strand hotspot of RelE (**Figure 2e**). In contrast, the five largest clusters of seed-guided designs were smaller, containing only 3–19% of the 493 high-confidence, high-contact structures (**Figure 2e**), and spanned a variety of binding topologies, including parallel and antiparallel β-strand pairing with the RelE β-strand.

From equal starting pools of 12,000 designs, seed-guided RFdiffusion retained substantially more structures at each filtering step: 1,786 designs (14.9%) maintained ≥ 26 contacts with RelE compared to only 173 (1.4%) from default RFdiffusion, and 493 (4.1%) versus 146 (1.2%) additionally satisfied ipAE < 10 (**Figure 2f**). Clustering these quality-filtered designs revealed that seed-guided generation produced 3.5-fold more structurally distinct clusters than RFdiffusion did, despite starting from an identical number of structures. Notably, the sharpest divergence between methods occurred at the contact filter, where seed-guided designs were more than 10-fold more likely to form extensive interfaces with the target, consistent with the structural priors provided by the seed motifs directing backbone generation toward productive binding geometries.

### Screening and Enrichment of Seed-Guided *De Novo* Protein Binders

To experimentally test designs generated by seed-guided RFdiffusion, we generated ∼52,000 candidates and applied the filtering pipeline in **Figure 1e**. Briefly, we assessed the compatibility of the designed sequences and structures using residue-level alignment (RLA) (25) and AF2 IG pLDDT and ipAE. We also selected designs for which the seed locations relative to RelE changed by no more than 10 Å when modeled using AF2 IG and for which the design was predicted to engage the RelE active site. Only 456 designs, representing 0.9% of all designs, met the filtering criteria. We used ProteinMPNN to redesign the sequences for these 456 backbones, retaining those for which AF2 IG predicted ≥ 3 contacts with the active-site residues of RelE, with contacts defined by an interchain distance of ≤ 3 Å, a feature that was observed for only one of the 12,000 RFdiffusion designs. A final set of 1,402 unique seed-guided sequences was chosen for experimental screening.

We screened designed binders for neutralization of RelE toxicity in *E. coli* in a survival-based assay (**Figure S6**) (19, 26). Neutralization activity was quantified by the enrichment of sequences in the library post-selection relative to pre-selection using the number of next-generation sequencing reads after two rounds of screening: enrichment ratio (ER) = log_2_[(frequency of reads after sorting)/(frequency of reads in the input library)]. Thirty-one designs were enriched relative to the input (**Figure 3a**), representing 2.2% of the library sequences. Among the 1,397 designs with enrichment scores, all of which had already passed stringent computational filters, the metrics used to score the designs before screening (i.e., RLA, ipAE, pLDDT) did not correlate with ER values (**Figure S7**). AF2 IG models for designs with ER > 1.0 all exhibited seed backbone atom RMSD < 3.3 Å relative to the originally docked seed positions (**Figure S8**).

**Figure 3.**
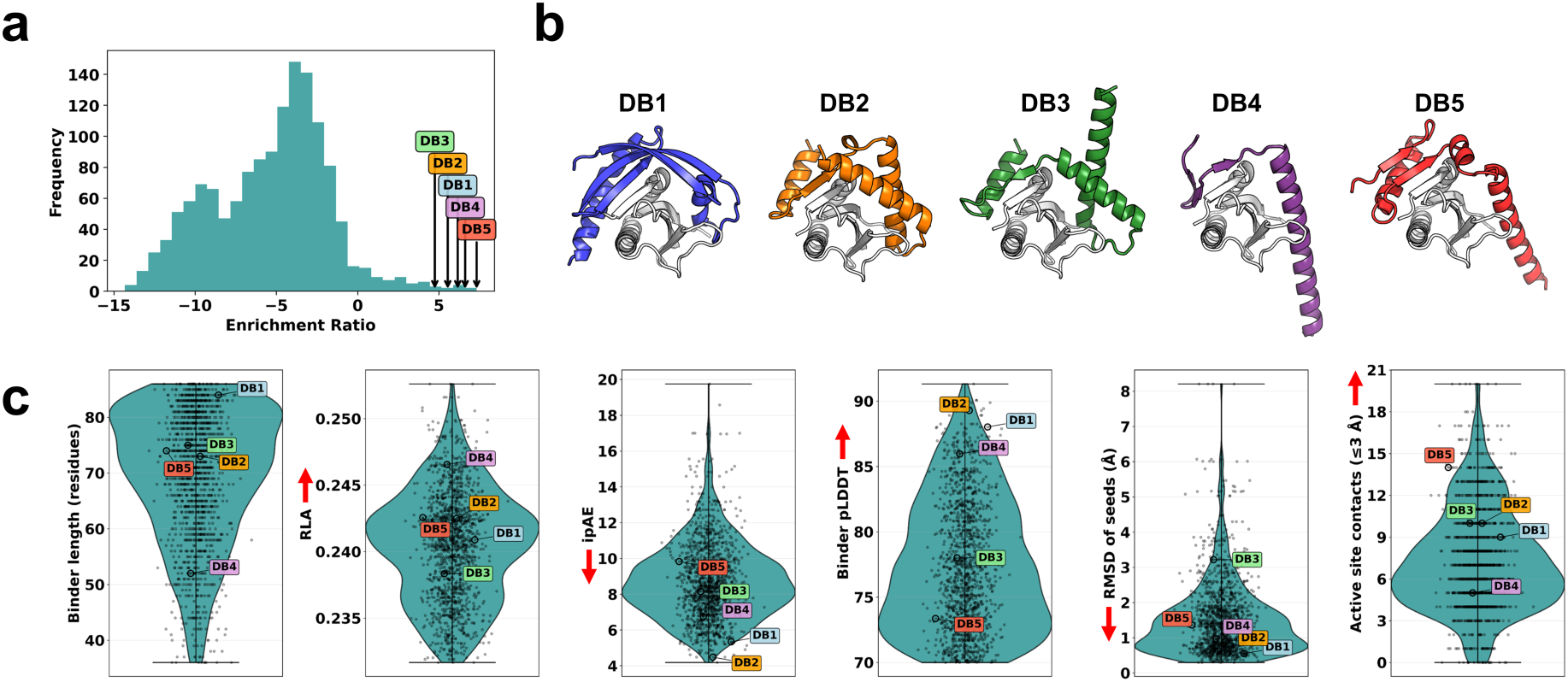
Experimentally screened seed-guided designs exhibited a range of enrichment ratios in the toxicity inhibition screen and scores for designed structures. (**a**) Distribution of enrichment ratios after two rounds of selection. (**b**) AF2 IG models for DB1-DB5. (**c**) Violin plots for the distribution of metrics used for filtering. RLA is the residue-level alignment score of Birnbaum et al. (25); RMSD of Seeds is the RMSD between the backbone atoms of the initially placed seed residues and their final positions after AF2 IG modeling; Active-site contacts reflect the number of atoms within 3 Å of active site residues. Red arrows indicate the desired direction for the filtering metrics, and scores for DB1-DB5 are indicated.

### Analysis of Seed-Guided *De Novo* Binder Hits

We selected five designed binders, DB1–DB5 (**Table S1**), for further characterization based on their high enrichment ratios and structural diversity and to span a range of seed RMSD, ipAE, and pLDDT scores (**Figure 3b, c**). The designed structures exhibited varied secondary structure compositions calculated by STRIDE (**Figure S9**), similar to the broader seed-guided design population. The rank order of the designs by enrichment ratio was DB5, DB4, DB1, DB2, DB3, with enrichment ratios of 7.3, 6.6, 6.2, 5.5, and 4.8, respectively (**Figure 3a, Table S2**). We confirmed that DB1–DB5 rescued cell growth in assays with RelE under control of the P_bad_ promoter when tested individually (**Figure S10**). These five designs were then tested for their ability to rescue cell growth using a toxicity spotting assay that is more stringent than the pooled screen, with RelE under the control of a vanillate-inducible promoter (P_van_). We quantified rescue using an inhibition ratio, defined in the Methods. Under these conditions, only DB1 and DB2 rescued cell growth with an inhibition ratio ≥ −4, although these designs did not have the highest enrichment ratios in the screen (**Figure 4a**).

**Figure 4.**
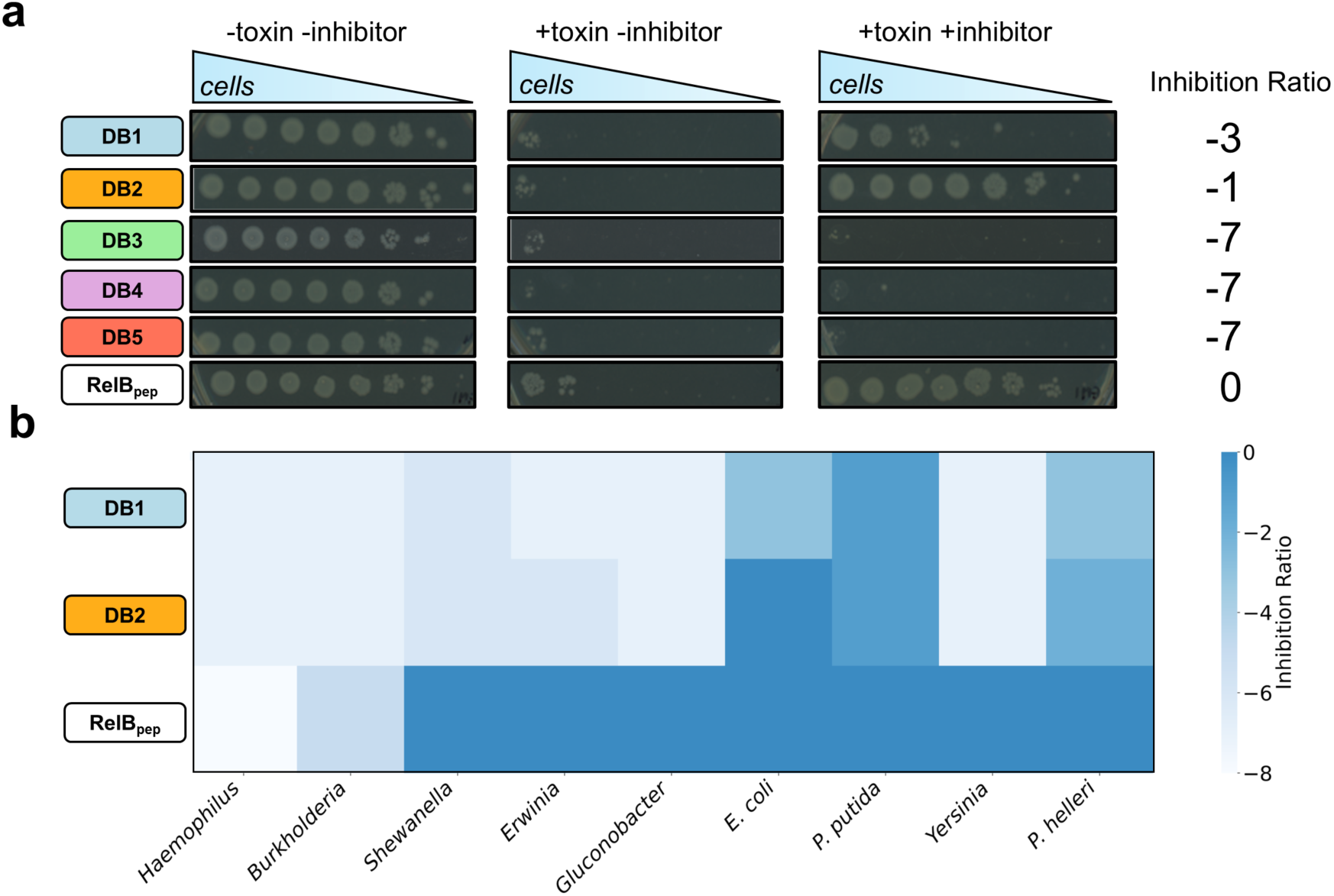
Functional analysis of designs DB1–DB5. (**a**) DB1, DB2, and control peptide RelB_pep_ inhibit RelE toxicity. Panels show growth of 10-fold dilutions of *E. coli* with or without expression of the RelE toxin and designed peptide or RelB_pep_ inhibitor. Inhibition ratio, as defined in the Methods, quantifies the effectiveness of designs in rescuing growth (a less negative ratio corresponds to better rescue). (**b**) Heatmap showing reduced growth rescue for DB1 and DB2 compared to RelB_pep_ in toxicity spotting assays using RelE orthologs from other bacterial species expressed in *E. coli* relative to uninduced controls.

Next, we tested whether DB1 and DB2 rescued the growth of *E. coli* expressing RelE orthologs from other bacterial species (19). RelB_pep_ provided cell rescue with an inhibition ratio ≥ −4 for seven of nine orthologous RelE toxins tested, with the two exceptions being toxins from *Haemophilus* and *Burkholderia*. In comparison, DB1 and DB2 provided cell rescue with an inhibition ratio ≥ −4 for only three of the nine toxins tested: *E. coli*, *P. helleri*, and *P. putida*, indicating higher specificity (**Figure 4b**).

We measured the binding of DB1–DB3 to RelE using biolayer interferometry (BLI) (**Table 1**, **Figure S11**). DB2 bound with an off-rate (k_off_) that was too slow to quantify in this assay; we estimated k_off_ < 10^-5^ s^-1^, bounding the dissociation constant at K_D_ < 0.7 nM. In contrast, DB1 and DB3 bound reversibly, and fitting yielded dissociation constants, K_D_ (k_off_/k_on_), of 30 nM and 6.5 μM, respectively (**Table 1**). RelBpep bound to RelE with a fitted dissociation constant of 2.1 nM, in reasonable agreement with previously reported surface plasmon resonance (SPR) measurements of full-length RelB binding to RelE (K_D_ = 0.3 nM) (26). This interaction was inhibited by increasing concentrations of DB2, which binds to RelE but not RelB_pep_, consistent with the two proteins binding to overlapping sites on RelE (**Figure S12**).

**Table 1.** Kinetics for designed binders and RelB_pep_ binding to RelE.

| Binder | RelE |  |  | RelE D6K |  |  | RelE K13E |  |  |
| --- | --- | --- | --- | --- | --- | --- | --- | --- | --- |
| | $k_{on}$<br>(M<sup>-1</sup> s<sup>-1</sup>) | $k_{off}$<br>(s<sup>-1</sup>) | $K_D$<br>(M) | $k_{on}$<br>(M<sup>-1</sup> s<sup>-1</sup>) | $k_{off}$<br>(s<sup>-1</sup>) | $K_D$<br>(M) | $k_{on}$<br>(M<sup>-1</sup> s<sup>-1</sup>) | $k_{off}$<br>(s<sup>-1</sup>) | $K_D$<br>(M) |
| DB1<sup>1</sup> | $6.2 \pm 0.0 \times 10^3$ | $1.8 \pm 0.2 \times 10^{-4}$ | $3.0 \pm 0.4 \times 10^{-8}$ | $3.7 \pm 0.1 \times 10^3$ | $9.4 \pm 0.5 \times 10^{-4}$ | $2.5 \pm 0.0 \times 10^{-7}$ | $3.6 \pm 0.1 \times 10^3$ | $4.5 \pm 1.7 \times 10^{-4}$ | $1.3 \pm 0.5 \times 10^{-7}$ |
| DB1 E63K<sup>2</sup> | $2.7 \pm 0.1 \times 10^3$ | $8.0 \pm 1.2 \times 10^{-4}$ | $3.0 \pm 0.3 \times 10^{-7}$ | $4.7 \pm 0.3 \times 10^3$ | $1.5 \pm 0.2 \times 10^{-3}$ | $3.2 \pm 0.3 \times 10^{-7}$ | $1.1 \pm 0.0 \times 10^4$ | $3.1 \pm 0.4 \times 10^{-4}$ | $2.9 \pm 0.5 \times 10^{-8}$ |
| DB2<sup>1</sup> | $1.4 \pm 0.0 \times 10^4$ | $< 1.0 \times 10^{-5}$ | $< 7.2 \pm 0.2 \times 10^{-10}$ | $3.0 \pm 0.0 \times 10^4$ | $2.1 \pm 0.3 \times 10^{-4}$ | $6.8 \pm 0.9 \times 10^{-9}$ | $8.3 \pm 0.0 \times 10^3$ | $1.1 \pm 0.0 \times 10^{-3}$ | $1.3 \pm 0.0 \times 10^{-7}$ |
| DB2 R20E<sup>2</sup> | $2.0 \pm 0.0 \times 10^4$ | $3.3 \pm 0.2 \times 10^{-4}$ | $1.6 \pm 0.0 \times 10^{-8}$ | $3.3 \pm 0.0 \times 10^4$ | $< 1.0 \times 10^{-5}$ | $< 3.1 \pm 0.0 \times 10^{-10}$ | $3.0 \pm 0.2 \times 10^3$ | $1.1 \pm 0.0 \times 10^{-3}$ | $3.8 \pm 0.2 \times 10^{-7}$ |
| DB3<sup>1</sup> | $8.0 \pm 7.2 \times 10^3$ | $3.2 \pm 0.6 \times 10^{-2}$ | $6.5 \pm 4.3 \times 10^{-6}$ | — | — | — | — | — | — |
| RelB<sub>pep</sub><sup>1</sup> | $3.4 \pm 1.2 \times 10^4$ | $7.2 \pm 3.5 \times 10^{-5}$ | $2.1 \pm 0.3 \times 10^{-9}$ | $9.9 \pm 1.0 \times 10^4$ | $< 1.0 \times 10^{-5}$ | $1.0 \pm 0.1 \times 10^{-10}$ | $1.4 \pm 0.2 \times 10^5$ | $1.2 \pm 0.6 \times 10^{-4}$ | $8.9 \pm 5.4 \times 10^{-10}$ |

<sup>1</sup> Determined from three independent replicates at seven analyte concentrations.
<sup>2</sup> Determined from two independent replicates at three analyte concentrations.
Values preceded by < indicate that koff is below the BLI instrument detection limit ( $1 \times 10^{-5}$ s<sup>-1</sup>); corresponding $K_D$ values are reported as upper bounds.
— not determined (DB3 was not assayed against RelE charge-swap variants).

### Designs Exhibit Context-Dependent Structures

Our seed-guided binder design protocol generated structures predicted to make many contacts with RelE. To explore whether these sequences were likely to fold to these structures in the absence of RelE, we used AlphaFold3 to predict the unbound conformations of 1,402 filtered designs selected for experimental screening (27). AlphaFold3 predicted substantial differences between isolated and RelE-bound conformations for many designs; the average RMSD between predicted structures of the bound and unbound states of the designs was 9.0 ± 4.1 Å (**Figure 5a**). Most designs (∼70%) showed higher pLDDT in the bound conformation (**Figure 5b**). For example, DB1 and DB3 exhibited pLDDT scores < 70 in the unbound conformation, but pLDDT > 70 when modeled in the bound conformation with AlphaFold3. AlphaFold3 predicted that DB1 and DB2 would undergo significant conformational changes upon binding, with unbound-to-bound RMSD of 5.8 Å and 10.2 Å, respectively (**Figure 5a**).

**Figure 5.**
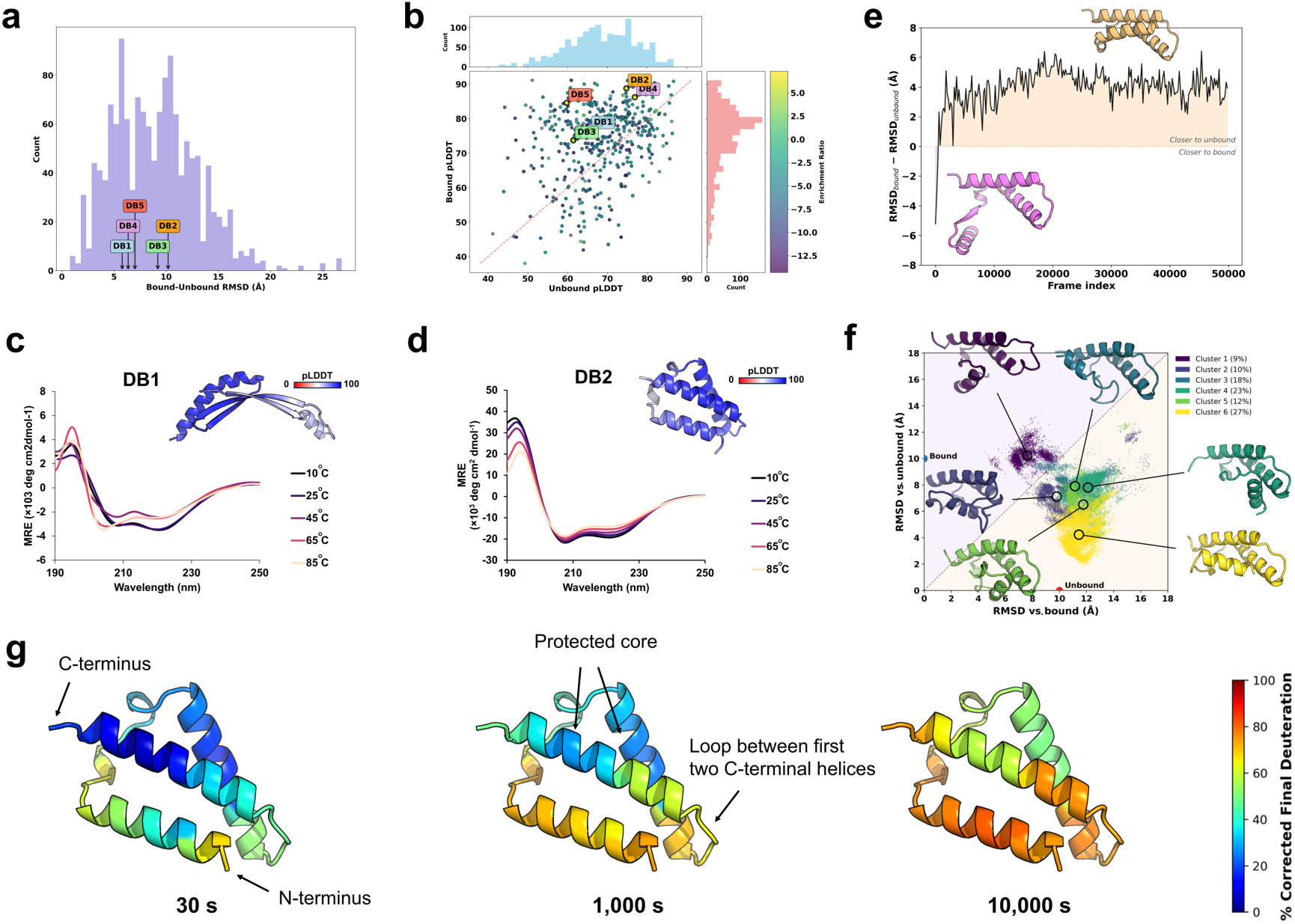
Designed binders exhibit context-dependent structures. (**a**) AlphaFold3-predicted differences in structure, by backbone Cα RMSD, between the bound and unbound structures for final filtered designs. (**b**) AlphaFold3 pLDDT confidence scores for predicted bound and unbound structures. (**c**,**d**) CD wavelength scans at increasing temperatures for (**c**) DB1 and (**d**) DB2, revealing high thermal stability of helical structure in unbound DB2. Inset: AlphaFold3-predicted structures of unbound DB1 and DB2, colored by pLDDT. **(e)** REMD simulation of DB2 shows a transition from the predicted bound conformation (pink) to structures more closely resembling the predicted unbound conformation (orange). Block-averaged trajectory (from 200 consecutive frames) RMSD_bound_ − RMSD_unbound_ as a function of simulation time. (**f**) Structures from the REMD simulation in panel e were projected based on their RMSD relative to the bound and unbound AlphaFold3 reference structures. Six clusters identified by k-means clustering on backbone Cα coordinates are colored by a gradient from most bound-like (Cluster 1, dark) to most unbound-like (Cluster 6, bright). Representative structures corresponding to the centroid for each cluster are shown, illustrating a structural transition from the bound to the unbound fold. Red and blue dots indicate the positions of the unbound and bound AlphaFold3 reference structures, respectively. Orange and purple shading denote unbound-favored and bound-favored regions, respectively. (**g**) Ribbon structures of DB2 are colored by fractional D uptake at three time points. Data are not normalized to FD controls.

We measured the structure and thermostability of unbound DB1 and DB2 using circular dichroism (CD) spectroscopy. CD wavelength scans at 25 °C revealed primarily β-sheet structure for DB1 and helical structure for DB2 (**Figures 5c and 5d, Table S3**) (28). DB2 exhibited minimal decrease in mean residue ellipticity (MRE) up to 85 °C (**Figure 5d**) and eluted as expected for a monomer in size-exclusion chromatography (**Figures S13b and S13c**). In contrast, DB1 had a lower melting temperature (T_m_) of 34 ± 2 °C (**Figure S14**) and was minimally soluble, precipitating at concentrations above ∼30 μM. DB1 injected at ∼10 μM eluted primarily as a large aggregate in size-exclusion chromatography (**Figure S13a**, **c**). Using mean residue ellipticity to estimate secondary structure, and assuming that the 35 residues of the N-terminal purification tag on DB2 do not contribute to the helical or strand content, 80%, 75%, and 62% of DB2 residues are estimated to be helical at 10, 25, and 65 °C, respectively (28). At 25 °C, this helical content is consistent with the AlphaFold3-predicted structure for unbound DB2, which is 78% helical (28).

To explore what structures DB1 and DB2 are likely to adopt in the absence of RelE, we performed REMD simulations initiated from the AF3-predicted bound conformation after removing the RelE chain (29–32). For DB2, the structure underwent a rapid change over the first 5 ns of the demultiplexed 300 K trajectory, with the 25 N-terminal residues exhibiting high conformational heterogeneity (RMSF up to ∼13 Å). In contrast, residues 26–73 adopted a relatively well-structured helical core, similar to the predicted bound structure (RMSF of ∼2–5 Å; **Figure S15**). In the DB1 simulation, the structure exhibited substantial conformational heterogeneity across the protein (**Figure S16a**) and sampled an ensemble of structures dissimilar from both the AlphaFold3-predicted bound and unbound conformations (**Figures S16b and S16c**), consistent with its low thermal stability (**Figure 5c**, **Figure S14**).

A closer analysis of the DB2 REMD simulation revealed that the N-terminal (residues 1–25) and C-terminal core (residues 26–73) largely retain their local folds (**Figures S17a and S17b**) but undergo significant motion relative to one another (**Figures S17c and S17d**). We aligned structures from the simulation using the less-variable C-terminal core and computed RMSD values for all residues with respect to the AlphaFold3-predicted bound and unbound reference models. The backbone reaction coordinate RMSD_bound_ − RMSD_unbound_, plotted in **Figure 5e**, highlights how the ensemble rapidly departed from the AlphaFold3-predicted bound conformation and became more similar to the AlphaFold3-predicted unbound conformation (RMSD_bound_ − RMSD_unbound_ > 0). **Figure 5f** illustrates the range of sampled structures using a two-dimensional RMSD landscape after C-terminal core alignment. K-means clustering of the ensemble resolved six states, of which the dominant clusters (Clusters 2–6, 91%) were more similar to the predicted unbound structure, with only Cluster 1 (9%) sampling bound-like conformations. A similar analysis for the N-and C-terminal regions is included in **Figures S17e and S17f**. Together, these analyses predict that isolated DB2 does not maintain the predicted RelE-bound arrangement; rather, the N-terminal region is dynamic relative to a more stable pair of C-terminal helices. Secondary structure analysis of 10,000 REMD frames sampled across all clusters yielded an average helical content of 61% for DB2, lower than the CD-derived estimate of 75% at 25 °C that may have been affected by the appended N-terminal tag.

To compare the predictions of the REMD simulations with the properties of DB2 in solution, we used hydrogen-deuterium exchange monitored by mass spectrometry (nHDX-MS) (33). To slow exchange kinetics and capture meaningful dynamics across highly flexible regions, labeling was performed at 4 °C, and we obtained excellent peptide coverage (**Figure S18, Dataset 1**). The experiments used DB2 with a 35-residue N-terminal tag (**Table S1**), but for consistency with the REMD analysis, here we refer to the first residue of DB2 as position 1. nHDX-MS showed that residues 1–25 were highly dynamic, with residues 1– 4 incorporating ∼90–94% D at 30 s. Residues 5–21 showed partial protection at 30 s, with most exceeding 80% by 1,000 s (**Figures 5g** and **S19a,b**). This result is consistent with the high backbone RMSF observed for this region in the REMD simulations. In contrast, the C-terminal helical core (DB2 positions 26–73) was substantially protected, with the two helices displaying mean D-uptake of ∼22% and ∼10% at 30 s, consistent with REMD RMSF values for residues in this region that are below 2.5 Å (**Figures 5g, S15, and S19b**). The most protected segment of the entire protein, centered on positions 61–67, incorporated only ∼7–10% deuterium at 30 s, confirming the presence of a well-ordered helical core that persists in the DB2 monomer and is consistent with secondary structure observed at high temperatures by CD (**Figure S19a,b**). The intermediate protection of the loop between the two C-terminal helices (∼56% D at 30 s; DB2 positions 42–60) and fast deuteration (∼72% at 300 s and 82% at 1,000 s) mirrors the local flexibility captured in simulation (mean RMSF ∼3.2 Å). Quantitative comparison of the FD-normalized single-residue D-uptake with the per-residue RMSF from REMD yielded a correlation that increased with labeling time (Spearman r = 0.73 at 30 s to r = 0.84 at 10,000 s), demonstrating that REMD captures regional differences in flexibility observed by nHDX-MS (**Figure S19c**). Because nHDX-MS was performed on tagged DB2, whereas REMD used untagged DB2, we use caution in interpreting the protection and flexibility near the N terminus. Together, the nHDX-MS data and REMD simulations establish a coherent picture of DB2 in the unbound state, with a thermostable C-terminal helical scaffold and a dynamic N-terminal arm.

### Investigating Predicted Binding Modes of DB2 by Mutational Analysis

To test whether DB2 binding to RelE is consistent with the designed binding mode, we used a mutational analysis focused on interchain salt bridges. RelE residue D6 is in a β-strand that is predicted to pair with a parallel strand in DB2, forming a salt bridge with DB2 R20 (**Figures 6a and 6b**). We generated the RelE D6K mutant and the compensatory DB2 R20E mutant and used BLI to quantify binding affinities. Mutation D6K in RelE weakened binding to DB2, from K_D_ < 0.7 nM for native RelE to 6.8 nM for the mutant, and mutation R20E in DB2, which we predicted by structural modeling to re-form the salt bridge, restored binding to RelE D6K (K_D_ < 0.3 nM) (**Figure 6b**, **Table 1**). We performed similar charge-swap mutation experiments using RelE K13E and DB1 mutation E63K (**Figure S20a**). BLI assays showed diminished and rescued binding affinities for complexes that respectively disrupted (RelE K13E with DB1) and restored (RelE K13E with DB1 E63K) the predicted interfacial salt bridge (**Figure S20b, Table 1**).

**Figure 6.**
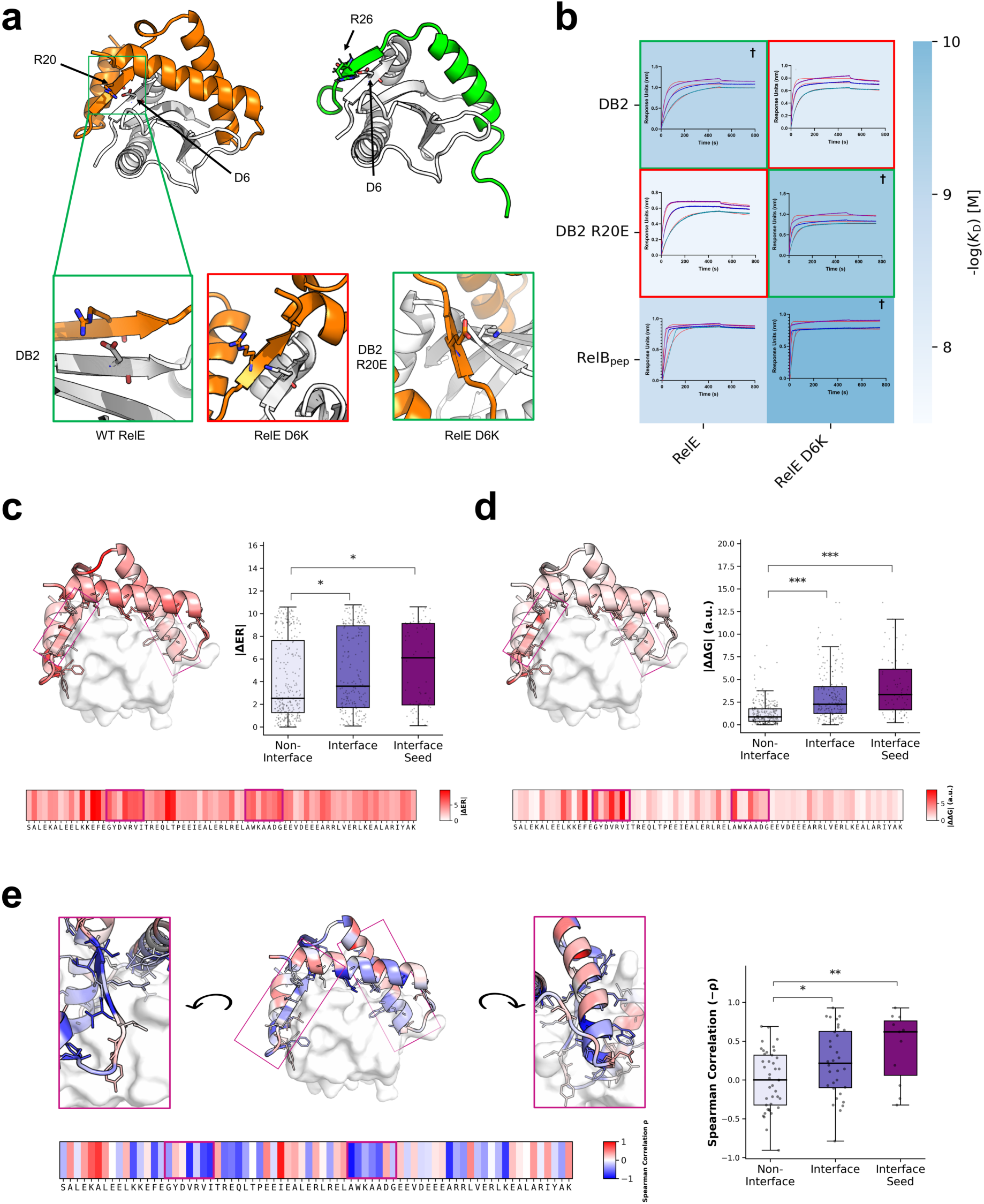
Mutational effects are consistent with the designed binding mode of DB2. **(a)** AlphaFold3-predicted structure of DB2 bound to RelE, and crystal structure of RelB_pep_ bound to RelE (PDB: 2KC8), indicating sites of charge-swap mutations. Boxes show modeled structures for mutants with oppositely charged residue pairs (green) or like-charged residue pairs (red). (**b**) Heatmap of average −log(KD) measured by BLI, with red and green boxes highlighting charge pairs as in panel a. Binding affinity is reported as the average of two independent replicates. Representative binding curves and fits are shown within the corresponding matrix cells. † indicates a reaction with immeasurably slow dissociation kinetics, bounded by k_off_ < 10^-5^ s^-1^. (**c**–**e**) Results of the mutational scanning experiment for DB2. (**c**) Averages of the impacts of six or seven mutations per position on the enrichment ratio, reported as average |ΔER|, mapped onto the AlphaFold3-predicted structure of DB2 in complex with RelE. Magenta boxes highlight residues from the seed regions. At right are distributions of |ΔER| values for mutations at non-interface positions, interface positions, and interface seed positions, as defined in the Supplementary Methods. (**d**) Average |ΔΔG_bind_| values from PottsMPNN as a per-position heatmap and mapped onto the AlphaFold3-predicted structure of DB2 in complex with RelE. Magenta boxes and box plots as in panel c. (**e**) Per-position Spearman correlation between ΔER and ΔΔG_bind_ for mutations, mapped onto the AlphaFold3-predicted structure of DB2 in complex with RelE. Magenta boxes and box plots as for panels **c** and **d**. * p-value < 0.05; ** p-value < 0.01; *** p-value < 0.001 by a Mann-Whitney U test.

Extending our mutational analysis, we tested point mutations at all positions of DB2 using a variation of the survival-based assay in which RelE was controlled by a vanillate-inducible promoter (P_van_) (**Figure S21a**). Substitutions to A, D, K, F, L, S, and V tested the effects of diverse hydrophobic and charge substitutions on function (34). Deep sequencing revealed an average absolute change in enrichment ratio compared to DB2 (|ΔER|) of 4.4 (SEM = 0.2, n = 476). However, the mutational scan was constrained by its dynamic range: 125 of 476 variants showed complete loss of function and were all assigned a pseudocount of 1, potentially underestimating the impact of some mutations. Nevertheless, we analyzed |ΔER| values by position to test whether residues that were predicted by AlphaFold to engage RelE gave larger effects (**Figure 6c**). We hypothesized that residues that are part of the interface region – i.e., residues of DB2 within 4 Å of RelE heavy atoms – would have lower mutational tolerance. Indeed, the 32 interface residues had an average |ΔER| of 4.8 (SEM = 0.3, n = 204), significantly higher than the average |ΔER| of the 41 non-interface residues, which was 4.1 (SEM = 0.2, n = 272; **Figure 6c**). Notably, 11 of 14 residues from the part of DB2 defined by the original seeds fall at the binding interface. Mutations at these interface seed residues had an average |ΔER| of 5.4 (SEM = 0.4, n = 70), significantly higher than that for non-interface residues (**Figure 6c**).

We used PottsMPNN, a computational tool that predicts changes in binding energies (ΔΔG_bind_) given a backbone structure, to assess whether the experimental screening data were consistent with the structure of the designed interface (35). Because the screening data reflect changes in protein function (including, for example, changes in expression, solubility, and binding), we hypothesized that the data and the structure-based computational predictions would correlate best at the positions where the structure is most directly tied to binding function – i.e., at the binding interface. We computed ΔΔG_bind_ using PottsMPNN for the seven experimentally tested substitutions (|ΔΔGbind| = 2.1 a.u., SEM = 0.1 a.u., n = 476; **Figure S21b**). PottsMPNN predicted an average |ΔΔG_bind_| of 3.2 a.u. (SEM = 0.2 a.u., n = 204) at the interface, significantly higher than the predicted 1.3 a.u. (SEM = 0.1 a.u., n = 272) for residues outside the interface (**Figure 6d**). Consistent with changes in experimental |ΔER|, interface seed residues had an even higher average |ΔΔG_bind_| of 4.2 a.u. (SEM = 0.4 a.u., n = 70; **Figure 6d**). Comparing ΔΔG_bind_ predictions for the charge-swap mutation RelE D6K binding to DB2 and the compensatory charge-swap mutation DB2 R20E binding to RelE D6K revealed increased (3.0 a.u.) and decreased (-1.1 a.u.) ΔΔG values, respectively, consistent with experimental trends (**Figure 6b**, **Table 1**).

Across all positions, PottsMPNN binding energies correlated modestly with experimental enrichment ratios (Spearman correlation, ρ = −0.27). When analyzing mutational trends on a per-position basis, interface positions had a correlation of ρ = −0.22 (SEM = 0.08, n = 32), significantly stronger than that for non-interface positions, which had a correlation of ρ = 0.00 (SEM = 0.06, n = 41; **Figure 6e**). Moreover, interface residues that also lie in seeds showed a higher correlation of ρ = −0.42 (SEM = 0.14, n = 11; **Figure 6e**). The trends in |ΔER|, |ΔΔG_bind_|, and per-position correlations between ΔΔG_bind_ and ΔER were consistent for different definitions of interface residues.

## Discussion

Generative design methods have produced *de novo* binders for a variety of targets (5). However, the structural diversity of successful designs has been limited, with RFdiffusion showing predominantly compact α-helical bundles with restricted interaction interfaces. Lu *et al*. showed that the structural biases of different design methods (Chroma, Genie2, RFdiffusion, and Protpardelle) are distinct, although none capture the structural diversity of native proteins (4). Recently, models have been adapted to increase diversity. For example, Frank et al. reduced the helical content of designs by introducing a helical loss (36), and Jendrusch and Korbel showed that random secondary structure conditioning can be combined with the Salad generative model to increase the predicted structural diversity of designs (37). Conditioning designs on CATH protein fold classifications is another way to generate increased structural diversity (38), as is conditioning a generated scaffold to form a β-strand to complement the exposed β-edge in a target, as done by Sappington et al. (39).

Methodological bias that limits structural diversity, favoring highly designable folds that are typically compact and highly helical, makes it difficult to achieve high surface complementarity throughout an extended interface or to simultaneously engage multiple binding sites, as is observed for many native protein-protein interactions. In the system studied here, the antitoxin RelB wraps around the surface of the toxin RelE to engage two spatially distinct interfaces that include the active site (22). We found that neither default RFdiffusion (with the exception of a single design) nor BindCraft generated confident designs with this type of structure or generated designs that contacted the active site, even when directed to the active site with designated binding hotspots. We addressed this gap by using seed-guided motif scaffolding to anchor RFdiffusion backbone generation with PDB-derived tertiary motif fragments that complement the target surface. This interface-first approach yielded designs with substantially greater structural diversity and more extensive target contacts than unguided or hotspot-guided generative methods, without using information about the native interaction.

Experimental screening identified a small number of functional designs, ∼2% of those that were tested. Function in the toxicity-inhibition assay requires that proteins be well expressed and fold in the cell, both in the presence and absence of RelE, and act at the ribosome, where RelE cleaves mRNA. It is possible that designs that did not pass the functional screen could bind to RelE under different conditions. The success rate in this campaign may also reflect the demands of the unusual topologies of our designed binders, which do not resemble the compact helical bundles that have performed well in other work. Modeling with AlphaFold3 predicted that most of the binders designed in this campaign will undergo conformational changes upon binding to RelE. REMD simulations support this for DB1 and DB2, demonstrating that in the unbound state, DB2 is not stable in the predicted bound conformation and can adopt a more compact structure. Our data support a model in which the C-terminal 48 residues of DB2 form a two-helix core, and the N-terminal 25 residues have some helical content but are flexible and can transition between the RelE-bound conformation and alternate states that are predicted to partially mask the binding interface. nHDX-MS corroborated this picture, revealing a dynamic N-terminal arm and a protected C-terminal helical core in the isolated protein, with D-uptake correlating well with REMD-derived RMSF. Our design process considered only one state, the bound state. The predicted alternate conformations of the unbound protein are likely favored because they bury residues that are engaged in an extensive hydrophobic interface when bound to RelE. Evaluating the conformational landscape of unbound designs may be informative for selecting designs that can, like DB2, mask large hydrophobic patches that promote binding via conformational changes that preserve solubility and improve success rates.

Despite extensive efforts, we were not able to obtain a crystal structure or nHDX-MS data for the RelE•DB2 complex. However, mutational data support the conclusion that DB2 engages RelE using the computationally introduced seed regions, in a geometry distinct from the native RelB binding mode. Specifically, charge-swap mutagenesis in one of the seed-derived regions of the DB2-RelE interface demonstrated that disrupting a predicted interchain salt bridge (RelE D6K) diminished binding affinity, and that a compensatory mutation in DB2 (R20E) restored it. Our mutational scanning of DB2 further revealed that predicted interface residues, particularly those within the seed regions, exhibited significantly lower mutational tolerance than non-interface positions. PottsMPNN binding energies, computed using the predicted complex structure, recapitulated this hierarchy: |ΔΔG_bind_| values were highest at interface seed positions, and per-position correlations between predicted ΔΔG_bind_ and experimental enrichment improved progressively from non-interface residues to interface residues to interface residues located within seeds. The convergence between structure-based predictions and experimental data provides evidence that seed-guided RFdiffusion successfully nucleated an interface with a geometry that matches the computational model and that is distinct from the native RelB/RelE interaction. Although our designed binders contact the same sites engaged by RelB and use similar secondary-structure elements, the designed chain runs in the opposite orientation at these contacts and forms a very different overall structure.

DB1 and DB2 exhibited more restricted functional rescue across RelE orthologs than did RelB_pep_. Designs that have extensive shape complementarity to their binding partner, as generated through seed-guided design, can engage a large interface that is less likely than a more localized binding site to be highly conserved across divergent orthologs. We anticipate that designing high surface complementarity, which arises naturally from multi-site seed-guided scaffolding, can be leveraged to engineer selectivity. This may be valuable for applications such as selective toxin inhibition in bacteriophage therapy, where the goal is to target only a specific pathogenic bacterial strain (16–18). In the future, combining seed-guided design that generates extensive contacts with computational methods that explicitly disfavor off-target binding may provide even greater ortholog specificity.

Our results demonstrate that incorporating structural priors from the PDB into generative backbone design can yield high-confidence predictions of protein structures with extensive target contacts, and that these translate to functional binders. Gainza et al. (40) and Balbi et al. (41) used a different approach, MaSIF-seed, to dock protein fragments from the PDB to targets as a starting point for design. In a recent campaign to identify binding sites on human cell-surface proteins, they generated miniproteins that engage FGFR2, IFNAR2, and HER3 with sub-to low-micromolar dissociation constants and computationally predicted success rates comparable to those of hotspot-directed design using RFdiffusion (41). By leveraging the structural diversity present in the PDB, seed-guided design approaches address limitations of unconstrained generative methods, enabling rational targeting of challenging protein-protein interfaces without requiring knowledge of the native binding mode. Seed-guided scaffolding provides a systematic framework for designing binders to targets requiring high surface complementarity, multiple binding sites, or non-canonical interaction geometries, thereby expanding the scope of tractable targets for computational protein design.

## Materials and Methods

The Supplementary Materials and Methods include detailed descriptions of procedures summarized here.

### Computational binder design

The design process started with peptide backbone fragments that complement the surface of RelE (PDB: 4FXE), generated in prior work using FASST(12, 19, 42–44). We selected the top 100 seeds at two surface sites and enumerated all site-1/site-2 seed pairs for RFdiffusion motif scaffolding in the presence of RelE. For each input, multiple backbones were generated. Sequences were designed with ProteinMPNN (45) and then evaluated with AlphaFold2 Initial Guess (AF2 IG) (20). In parallel, unguided RFdiffusion (5) and BindCraft (23) designs were generated using RelE hotspot residues derived from the RelB interface (22). Candidate designs were filtered by RLA sequence-structure compatibility (25), AF2 IG ipAE and pLDDT, length constraints for synthesis, active-site contact requirements, and preservation of the seed placement after AF2 IG modeling, yielding a smaller set of backbones that were then redesigned to produce the final library.

### Functional screening and tests of inhibition of RelE toxicity

The designed peptide library was synthesized as an oligo pool, cloned into the pKVS45 expression vector under the control of an anhydrotetracycline (aTc)-inducible P_tet_ promoter, and transformed into an MG1655 screening strain carrying inducible chromosomally integrated P_bad_-RelE at the *amyA* locus. Cells were first grown under conditions repressing toxin expression, then induced to express the designed binder, and then challenged with RelE induction on plates containing M9 medium supplemented with arabinose; surviving colonies were recovered, plasmid DNA was isolated, and the pooled screen was repeated to generate independent post-selection libraries. Library composition before and after selection was quantified by NGS, and sequence-level enrichment was calculated relative to the input pool. A similar survival-based framework was used for the mutational scanning of DB2 variants to measure mutational tolerance and functional robustness, with RelE toxin expression under the control of the vanillate-inducible P_van_ promoter. For the follow-up toxicity spotting assay (19), designed binders were tested for their ability to neutralize RelE in two MG1655 backgrounds, one carrying chromosomally integrated RelE and the other expressing RelE from a plasmid. Overnight cultures were serially diluted and spotted onto agar under inducing or control conditions, with binder expression driven by aTc and toxin expression induced by arabinose or vanillate as appropriate; the assay readout was the extent of growth rescue relative to control plates. Rescue was scored on an ordinal scale from complete rescue to no rescue, providing a simple functional measure of RelE neutralization.

### Molecular dynamics simulations

The molecular dynamics simulations used AMBER24 (29) with the ff14SB force field (30) and explicit solvent. Starting from AlphaFold3-predicted bound structures, structures were minimized and equilibrated through restrained NPT and NVT phases before production REMD. Production used 48 replicas spanning 300–450 K, with exchanges attempted every 1 ps for 500,000 total attempts, giving 500 ns per replica and 24 μs aggregate sampling per system; trajectories were analyzed at 300 K using RMSD, k-means clustering, and STRIDE secondary-structure assignment (24).

### Biophysical characterization of binders

Selected binders and RelE constructs were expressed recombinantly in *E. coli*, purified by affinity chromatography, and analyzed by SDS-PAGE and concentration assay before downstream studies. Binding kinetics were measured by biolayer interferometry using biotinylated binders on streptavidin sensors and serial dilutions of RelE, with competition experiments against RelB peptide and model-based fitting to extract k_on_, k_off,_ and K_D_. Secondary structure content and thermal stability were assessed using circular dichroism spectroscopy. Analytical size-exclusion chromatography was used to test for oligomerization, and nHDX-MS was used to quantify local protection patterns as a function of time at 4 °C.

### Use of generative AI

ChatGPT 5.5 and Claude Opus 4.4–4.6 were used for critical feedback on the logic and clarity of the manuscript, text-editing suggestions, and analysis-code generation. All outputs were reviewed and verified by the authors, and no AI tool was used to generate primary data or draw scientific conclusions.

## Associated Content

Supplementary Materials and Methods

Figures

Tables S1–S3

Supplementary References

Dataset 1 – Peptide-level deuterium uptake for DB2

## Acknowledgements

Research reported in this publication was supported by the National Institute of General Medical Sciences of the National Institutes of Health under award number R35GM149227. The content herein is solely the responsibility of the authors and does not represent the official views of any funding organization. The authors acknowledge the MIT Office of Research Computing and Data for providing high-performance computing resources that contributed to the results reported in this paper. We thank the BioMicro Center at MIT for conducting next-generation sequencing; M. Black for standards for SEC analysis; and L. Guan and S. Kotelnikov for helpful discussions and suggestions throughout this work. We thank V. Sarpe at Trajan Scientific and Medical for support with HDExaminer™ PRO.

