## Supplementary Information for "Seed-Guided *De Novo* Design Expands the Structural Diversity of Antitoxin Protein Binders"

### Supporting Information

### Supplementary Materials and Methods

**Materials.** Chemically competent BL21(DE3) *E. coli* cells, restriction enzymes, NEBuilder HiFi DNA Assembly Cloning Kit, KLD Enzyme Mix, OneTaq Quick-Load 2× Master Mix, and chemically competent DH5α cells were purchased from New England Biolabs. The Pierce bicinchoninic acid (BCA) assay kit and electrocompetent MegaX DH10B T1R cells were acquired from Thermo Fisher Scientific. KAPA HiFi HotStart ReadyMix was purchased from Roche. Chemically competent Rosetta (DE3) cells were purchased from Sigma-Aldrich. Octet streptavidin (SA) biosensors were purchased from Sartorius.

**Bacterial Strains and Plasmids.** The base strain for the study was *E. coli* strain MG1655. The strain used for high-throughput screening had a copy of $P_{bad}$-RelE integrated into the *amyA* locus (1). The peptide library was cloned into the pKVS45 vector under $P_{tet}$ control. To test inhibition of RelE orthologs, a low-copy pSC101/kanR plasmid expressing RelE variants was used under the control of an inducible Pvan promoter.

**Computational Design of De Novo Binders.** In prior work, seeds were generated around the target RelE protein (PDB: 4FXE ) using FASST (2) to identify surface-complementary peptide backbones defined by tertiary motifs (TERMs) in the PDB (3-5) and then ranked using a combination of scores that assessed RelE complementarity, as described by Ghose *et al.* (6). This work used the 100 highest-scoring seeds, which were located on either of two opposing sides of the RelE protein, denoted as site 1 and site 2, respectively. The seeds were divided into two clusters of 53 and 47 seeds based on their location. For every combination of one site-1 and one site-2 seed (a total of 4,982 unique seed pairs), seed backbone coordinates were combined into a single input PDB file. Each input file was used as a motif for the RFdiffusion motif-scaffolding protocol (7), in the presence of the RelE target protein, allowing 0–40 residues to be built at each of three locations: the N-terminus, between the seeds, and at the C-terminus. Five to six backbones were generated per input file. The protocol was repeated using combinations of Base_ckpt.pt and the Complex_beta_ckpt.pt model weights. Following backbone generation, ProteinMPNN (8) was used to assign a sequence. The designed complex was then evaluated using AlphaFold2 Initial Guess (9), resulting in a total of 51,692 structure predictions. Separately, RFdiffusion was used to generate 12,000 binders of RelE without any motif input, with lengths 44–144 residues, using Base_ckpt.pt model weights and using the following RelE hotspot residues: L5, D6, F7, D8, E9, R10, E40, A41, N42, K43, L44, R45, G46, E82, and A83. These residues were chosen because they form atomic contacts within 4 Å of RelB in PDB structure 2KC8 (10). We also tested BindCraft (11), using the same hotspot residues and the default_4stage_multimer settings. Because of the time required to produce designs passing the BindCraft filters, we generated only ~500 designs in 8.1 GPU-days (NVIDIA L40S), which may not fully represent the capabilities of the method when allocated a greater resource budget.

**Computational Filtering of De Novo Binders**. Seed-guided RFdiffusion designs were filtered using computed metrics. We identified the top 75% of designs by RLA score (12), which evaluates sequence-structure compatibility, and then chose designs with AlphaFold2 interface predicted aligned error (ipAE) < 20 and predicted local distance difference test (pLDDT) > 70. Designs with ≥ 87 residues were removed to facilitate DNA synthesis (Twist Biosciences). We also required a minimum of three contacts, defined as any pair of atoms within 3 Å, between the design and the active-site residues of RelE: R45, K52, and R61 (10). Finally, after aligning the backbone atoms of the RelE chain from RFdiffusion designs with the RelE chain after structure prediction by AlphaFold2, we required that the RMSD of the backbone atoms corresponding to the seeds be < 10 Å. After filtering, 456 designs remained. We then used ProteinMPNN to design five new sequences for each of these 456 backbones, and the new designs were included in the library if a minimum of three contacts within 3 Å of the active-site residues of RelE was preserved in the AlphaFold2 Initial Guess-predicted complex structure, resulting in 1,402 designed sequences.

**Structural Clustering Analysis of Designed Binders.** For structural diversity analysis, designs were first filtered by predicted interface quality, retaining only structures with ipAE < 10. The binder chain (chain A) was extracted from each AlphaFold2-evaluated complex. Structural clustering was performed using Foldseek (13) easy-cluster with the following parameters: minimum sequence identity of 0.0 (--min-seq-id 0.0) to rely exclusively on structural similarity, alignment coverage threshold of 0.8 (-c 0.8), bidirectional coverage mode (--cov-mode 0), TM-score-based alignment (--alignment-type 1), connected-component clustering (--cluster-mode 1), and a TM-score threshold of 0.5 (--tmscore-threshold 0.5). Independently, exhaustive all-versus-all pairwise TM-scores were computed using Foldseek structural alignment with TM-align (--exhaustive-search 1, -e inf, --alignment-type 1) for all binder chains in each dataset.

**AlphaFold3 Structure Prediction of Bound and Unbound Conformations.** We predicted the structures of designs alone or in complex with RelE using AlphaFold3 with default settings without templates and only the unpaired multiple sequence alignment of RelE (14). Interface residues of DB2 were identified from the AlphaFold3-predicted complex structure as those with any heavy atom within 4 Å of a RelE heavy atom.

**Designed Binder Library Assembly.** An oligonucleotide pool encoding each peptide plus 21 bp flanking sequences and BsmBI sites was ordered from Twist Biosciences, amplified, and cloned into the pKVS45 vector backbone under the control of an anhydrotetracycline (aTc)-inducible $P_{tet}$ promoter. Invitrogen MegaX DH10B *E. coli* cells were transformed by electroporation and recovered in 2×YT medium containing 200 µg/mL ampicillin, and the DNA library was purified by miniprep.

**High-Throughput Survival-Based Screening.** Designs were tested for their ability to prevent cell growth arrest upon induction of RelE, as established previously (6). Briefly, PCR products were electroporated into *E. coli* MG1655, which had a copy of $P_{bad}$-RelE integrated into the *amyA* locus, and recovered in SOC at

37 °C for 1 h before inoculation into 20 mL of M9 medium containing 35 µg/mL kanamycin, 200 µg/mL ampicillin, and 0.8% glucose to suppress expression of RelE. Following 30 min of incubation, designed binder protein expression was induced with 0.1 ng/µL anhydrotetracycline (aTc), followed by incubation at 37 °C for 30 min. Cells were pelleted and plated onto M9 medium containing 0.004% arabinose to express RelE and 0.1 ng/µL aTc. Plates were incubated overnight at 37 °C in the dark. Colonies were scraped and miniprepped to generate a post-selection plasmid library. The screen was repeated to generate a second post-selection library.

**Deep Mutational Scanning of DB2.** To assess the mutational tolerance of DB2, we performed a survival-based screen using *E. coli* MG1655 strains with a plasmid encoding the RelE toxin under the control of the vanillate-inducible Pvan promoter. An oligonucleotide pool encoding wild-type DB2 and DB2 variants with mutations at all positions to A, D, F, K, L, S, and V was ordered from Twist Biosciences to sample a range of hydrophobic, charged, and polar substitutions. The library was amplified, cloned into pKVS45 under aTc-inducible $P_{tet}$ control, and transformed into the screening strain as described above for the designed binder library. Following recovery in SOC, cells were grown in LB containing 35 µg/mL kanamycin and 200 µg/mL ampicillin. DB2 variant expression was induced with 0.1 ng/µL aTc for 30 min at 37 °C before cells were pelleted and plated onto LB agar containing 10 µM vanillate and 0.1 ng/µL aTc. Plates were incubated overnight at 37 °C in the dark, and plasmids were isolated from surviving colonies. The screen was repeated to generate a second post-selection library.

**Next-Generation Sequencing.** The naïve and post-selection libraries were prepared for next-generation sequencing (NGS) by amplification using KAPA HiFi, which appended Illumina sequencing adapters and barcodes as done previously (6). Final products were sequenced, generating up to 5 million reads on an Illumina MiSeq System. Variants that showed zero post-selection counts were assigned a pseudocount of 1. Sequences were normalized by the total number of reads for each sample to generate clone frequencies, and the enrichment ratio (ER) of each sequence was calculated using $\log_2$(frequency in the post-selection library/frequency in the input library) (6). Changes in enrichment ratios were calculated as the difference between the mutant and wild-type ER values ($\Delta ER = ER_{mutant} - ER_{wildtype}$).

**PottsMPNN Binding Energy Predictions.** Binding energies for DB2 mutations were calculated using a version of PottsMPNN (15) with weights fine-tuned to predict energies from the Mega-scale dataset (16). The top-ranked AlphaFold3-predicted structure (of five models by ranking_score) was used as input for PottsMPNN. To predict the bound binder conformation, we used AlphaFold3 with the binder sequence, the RelE sequence, and an unpaired RelE multiple sequence alignment. To predict the unbound structure for each design, we used only the binder sequence. For each mutation, the change in binding energy ($\Delta\Delta G_{bind}$) was computed as the difference of the mutant and wild-type binding energies of the bound and unbound states: $\Delta\Delta G_{bind} = \Delta\Delta G_{mutant} - \Delta\Delta G_{wildtype}$; $\Delta\Delta G_{mutant} = \Delta G_{mutant,bound} - \Delta G_{mutant,unbound}$; $\Delta\Delta G_{wildtype} = \Delta G_{wildtype,bound}$

– $\Delta G_{wildtype,unbound}$. Correlation of the predicted binding energies with experimental enrichment ratios from the mutagenesis screen was quantified using the Spearman correlation coefficient.

**Replica-Exchange Molecular Dynamics (REMD).** All simulations were performed using pmemd.cuda from AMBER24 (17) with the ff14SB protein force field (18), TIP3P (19) explicit solvent, a 9 Å nonbonded cutoff, and particle-mesh Ewald for long-range electrostatics. All molecular dynamics phases (NPT density equilibration, NVT equilibration, and REMD production) additionally used SHAKE constraints on bonds involving hydrogen and a Langevin thermostat ($\gamma$ = 2.0 $ps^{-1}$) for temperature control. System preparation (pdb4amber, tleap) and trajectory analysis (cpptraj) used AmberTools25 (20).

Starting structures were derived from AlphaFold3-predicted bound conformations, cleaned with pdb4amber, and parameterized in tleap. Each solute was solvated in a truncated octahedral box with a 10 Å buffer, neutralized with $Na^+$/$Cl^-$ counterions, and supplemented with NaCl to a target concentration of 0.15 M. Systems were energy-minimized for 3,000 cycles (1,500 steepest descent, 1,500 conjugate gradient) with positional restraints of 5.0 $kcal \cdot mol^{-1} \cdot Å^{-2}$ on non-hydrogen solute atoms. Box density was then equilibrated at 300 K using a segmented NPT protocol. The first segment (5 ps, 1 fs timestep, with 5.0 $kcal \cdot mol^{-1} \cdot Å^{-2}$ positional restraints on non-hydrogen solute atoms) used a Berendsen barostat for initial volume relaxation; all subsequent segments (20 ps each, 2 fs timestep, with restraints reduced to 2.0 $kcal \cdot mol^{-1} \cdot Å^{-2}$) used a Monte Carlo barostat. Segments were run iteratively until the relative change in mean box volume between consecutive segments fell below 0.5%. From the converged NPT configuration, 48 independent NVT equilibrations (200 ps each, 2 fs timestep) were performed at their respective target temperatures, with initial velocities drawn from a Maxwell–Boltzmann distribution.

Production temperature REMD used 48 replicas with temperatures distributed geometrically between 300 K and 450 K. Each replica was propagated in the NVT ensemble. Exchanges between neighboring replicas were attempted every 1.0 ps (500 MD steps; 2 fs timestep) for 500,000 total exchange attempts, yielding 500 ns per replica and 24 μs of aggregate sampling per system. Coordinates were saved every 10 ps, yielding 50,000 frames per replica. The continuous 300 K trajectory was reconstructed by demultiplexing the exchange log using cpptraj. Mean nearest-neighbor exchange acceptance probability ranged from 25–37%. Conformational analysis was performed on the 300 K trajectory using backbone RMSD relative to AlphaFold3-predicted bound ($RMSD_{bound}$) and unbound ($RMSD_{unbound}$) reference structures, with superposition computed using the full backbone for DB1 and the C-terminal region (residues 26–73) for DB2. K-means clustering (k = 7 for DB1 and k = 6 for DB2, selected by the elbow method) was performed on full-backbone Cα coordinates. Secondary structure from REMD was assigned for every fifth frame of the 300 K REMD replica (10,000 frames total) using STRIDE (21).

**Toxin Inhibition Assays.** RelE neutralization was tested using a toxicity spotting assay (6). pKVS45 plasmids encoding designed proteins were transformed into two *E. coli* MG1655 strain backgrounds: one carrying a chromosomally integrated copy of $P_{bad}$-RelE at the *amyA* locus, and one harboring a plasmid encoding RelE under $P_{van}$ control. Transformants were grown overnight at 37 °C in 2 mL LB supplemented with 35 μg/mL kanamycin and 200 μg/mL ampicillin, then serially diluted 10-fold (eight dilutions) and spotted onto plates containing the appropriate toxin inducer (0.01% arabinose, 0.1% arabinose, or 10 μM vanillate), aTc, both, or neither (control). Inducer-only plates confirmed that RelE expression was toxic. The difference in the number of dilutions yielding colonies between inducer-plus-aTc plates and control plates was calculated, yielding an integer rescue score from 0 (complete cell rescue, high RelE neutralization) to 8 (no rescue). The inhibition ratio is defined as:

$$inhibition\,ratio = \; log_{10}(cell\;growth + van + aTc) - \; log_{10}(cell\;growth - van - aTc)$$

where cell growth + van + aTc is the number of dilutions that supported colony growth on the plate with van and aTc, and cell growth − van − aTc is the corresponding number on the plate without van and without aTc.

**Protein Expression and Purification for Biophysical Characterization.** DNA sequences encoding selected designed binders were cloned into pET28a and pDW363 vectors using Gibson assembly; alternatively, constructs were purchased from GenScript or Twist Biosciences. Oligonucleotides encoding designed binder proteins or the RelB peptide ($RelB_{pep}$) sequence (K47–L79) (10) were cloned into pDW363 downstream of a sequence encoding SUMO and a hexahistidine tag and transformed into BL21 (DE3) cells. The RelE mutant constructs were cloned into pET28a and purchased from GenScript. The pET28a constructs were cloned with an N-terminal hexahistidine tag preceding the designed sequences. DB1 and RelE were cloned separately into the pET28a vector by Twist and included an additional N-terminal hexahistidine tag. Sequences for all constructs are given in **Table S1**.

Cloned pET28a constructs were transformed into Rosetta (DE3) cells, plated on LB agar containing 35 μg/mL kanamycin and 35 μg/mL chloramphenicol, and grown overnight at 37 °C. Colonies were selected and inoculated into 16 mL of LB and incubated overnight at 37 °C. Starter cultures were subsequently transferred into 800 mL of LB containing 35 μg/mL kanamycin and 35 μg/mL chloramphenicol and grown at 37 °C at 130 rpm in an Innova 4300 incubator shaker (New Brunswick Scientific) until the optical density at 600 nm ($OD_{600}$) reached approximately 0.6. Protein expression was induced with 200 μg/mL IPTG, followed by incubation for 4 h at 37 °C and 130 rpm for designed binders, or by incubation overnight at 37 °C and 130 rpm for RelE and RelE variants. Then, cells were harvested by centrifugation at 5,000 × g at 4 °C for 20 min in an Avanti J-E centrifuge (Beckman Coulter) and stored at −20 °C until purification. Expression was confirmed by SDS-PAGE analysis. Cells from resuspended pellets were lysed with a Q500 sonicator (Qsonica) using a 5-s-on, 5-s-off pulse sequence at 55% amplitude for a total pulse time of 2 min. The lysate was separated by centrifugation at 12,000 × g at 4 °C for 30 min in an Avanti J-E centrifuge (Beckman Coulter).

For designed proteins expressed from the pET28a vector, the supernatant was loaded onto a cobalt-charged HiTrap IMAC FF 5 mL column. Proteins were eluted with a gradient of imidazole from 0–500 mM in 50 mM Tris, 200 mM NaCl, pH 8.0. Purity was assessed using SDS-PAGE, and fractions containing pure protein were pooled and dialyzed in six consecutive 5 L volumes of phosphate-buffered saline (PBS, pH 7.4). The protein was concentrated using 3 kDa MWCO centrifugal filters, and its concentration was measured by BCA assay (Thermo Fisher Scientific).

RelE proteins were purified similarly, but under denaturing conditions using 8 M urea. Pure fractions of RelE protein were first dialyzed into one 5 L volume of PBS containing 1 mM dithiothreitol (DTT). Insoluble protein was separated by pelleting at 4,000 × g at 4 °C for 20 min and resolubilized in 50 mM Tris, 8 M urea, pH 8.0 buffer by adapting a protocol from the Strobel laboratory (22). Solubilized protein was dialyzed into one 5 L volume of 50 mM Tris, 70 mM $NH_4Cl$, 300 mM KCl, 7 mM $MgCl_2$, 1 M urea, and 1 mM DTT, pH 7.5, before concentration using a 10 kDa MWCO centrifugal filter. During concentration, the buffer was exchanged with a buffer containing 50 mM Tris, 70 mM $NH_4Cl$, 30 mM KCl, 7 mM $MgCl_2$, 1 mM DTT, and 20% glycerol at pH 7.5, followed by storage in the same buffer. Protein purity was assessed by SDS-PAGE. Protein concentration was measured by a reducing-agent compatible BCA assay (Thermo Fisher Scientific).

To prepare biotinylated proteins, designed proteins cloned into pDW363 were transformed into BL21 (DE3) cells, plated on LB agar plates containing 35 µg/mL kanamycin and 35 µg/mL chloramphenicol, and grown overnight at 37 °C. Colonies were used to inoculate 5 mL of LB containing 200 µg/mL ampicillin. Subsequently, 500 µL of overnight culture was used to inoculate 20 mL of LB medium containing 200 µg/mL ampicillin and 50 µM biotin. Cells were incubated until the $OD_{600}$ reached ~0.6 and were then induced with 200 µg/mL IPTG and incubated for another 5 h. Cells were harvested at 4,000 × g at 4 °C for 20 min before storage at −20 °C until purification; expression was confirmed by SDS-PAGE. Cells were thawed and lysed using 4 mL of B-PER reagent (Thermo Scientific) before shaking for 10 min at room temperature. A 1 mL sample of the lysate was then centrifuged at 15,000 rpm for 10 min, and the supernatant was incubated with 500 µL of Ni-NTA agarose resin (Invitrogen) equilibrated with 50 mM Tris, 200 mM NaCl, pH 8.0, washed three times with this buffer, and then eluted with 2 mL of 50 mM Tris, 200 mM NaCl, 500 mM imidazole, pH 8.0. The eluted protein was then stored at −20 °C. Purification was confirmed by SDS-PAGE.

**Protein Binding by Biolayer Interferometry (BLI).** Biotinylated protein was added at 2% (v/v) to a nonspecific-binding (NSB) buffer composed of PBS with 1% BSA, 0.01% Tween, 1 mM DTT, and 600 mM sucrose (23). RelE protein was serially diluted using seven 2-fold dilutions in NSB buffer from 2,000 nM to 30 nM with a reference well of NSB buffer only. An Octet R8 biolayer-interferometry instrument (Sartorius) was loaded with eight streptavidin-coated biosensors. Baseline signal was recorded in NSB buffer for 60 s, followed by loading of biotinylated designed binders or $RelB_{pep}$ until a 1-nm binding-response threshold was attained for all biosensors. Following 120 s in NSB buffer, protein-loaded biosensors were incubated with the serial dilutions of RelE for 500 s and then in NSB buffer for 500 s. Nonspecific binding was assessed by repeating the experiment using a biotinylated SUMO-His protein as a negative control. Experiments were repeated in triplicate using independent protein preparations of the biotinylated protein binder or $RelB_{pep}$. BLI experiments for charge-swap mutants were performed using three 3-fold dilutions in NSB buffer at 2,000 nM, 700 nM, and 200 nM under the same conditions, except dissociation was monitored for 200 s and assays were performed in duplicate using independent preparations of protein binders. Binding kinetics were fit to a 1:1 binding model using the association and dissociation kinetics equation in Prism 10 (GraphPad) with Bmax > 0 and a single value of kon > 0 shared across concentrations. A single value of koff was shared across concentrations and constrained to be > $10^{-5}$ $s^{-1}$. Slower dissociation could not be accurately quantified (24). Therefore, for binding reactions with fitted koff values of $10^{-5}$ $s^{-1}$ or lower when unconstrained, we report only an upper bound on the dissociation rate and the dissociation constant.

Statistical comparison of binding kinetics was performed using 95% confidence interval (CI) overlap analysis. Confidence intervals were calculated using profile likelihood in Prism 10 (GraphPad). Datasets were considered kinetically distinct if CIs for $k_{on}$ or $k_{off}$ did not overlap (equivalent to $p < 0.05$). $K_D$ confidence intervals were propagated from $k_{on}$ and $k_{off}$ using $K_{D,\text{lower}} = k_{\text{off,lower}}/k_{\text{on,upper}}$ and $K_{D,\text{upper}} = k_{\text{off,upper}}/k_{\text{on,lower}}$.

We tested for competition between DB2 and $RelB_{pep}$ binding by loading streptavidin-coated biosensors with biotinylated $RelB_{pep}$ followed by association with 3-fold serial dilutions of DB2 from 2,000 nM to 8 nM in the presence or absence of 500 nM RelE. This was repeated in duplicate using independent protein preparations of DB2 and $RelB_{pep}$. Binding kinetics were fit to a 1:1 binding model, implemented in Prism 10 (GraphPad) using the “Association kinetics—two or more concentrations of hot ligand” equation. $B_{max}$ values were used to determine equilibrium bound and unbound response values.

**Circular Dichroism (CD) Spectroscopy.** Wavelength scans were performed on a Jasco J-1500 spectrometer from 190 to 250 nm in 1 nm steps, using tagged constructs pET28a/DB1 and pET28a/DB2 at 10–15 μM in PBS buffer, pH 7.4. For DB2, scans were collected for three independent protein preparations at 10, 25, 45, and 65 °C and for two preparations at 85 °C. For DB1, scans were collected in technical triplicate at all temperatures. Temperature scans were performed from 10 °C to 95 °C, at 1 °C/min, in duplicate. The mean residue ellipticity (MRE) was calculated using established methods (25). The BeStSel algorithm was used to estimate secondary structure content (26).

**Analytical Size-Exclusion Chromatography.** 500 μL samples of 400 μM DB2 (pET28a/DB2) and 10 μM DB1 (prepared from pET28a/DB1) were injected into a fast protein liquid chromatography (FPLC, AKTA pure, GE Healthcare) system equipped with a Superdex 200 10/300 GL Size Exclusion Chromatography (SEC) column and eluted with buffer containing 20 mM Tris and 300 mM NaCl at pH 8.0 using a flow rate of 0.4 mL/min at 4 °C. Protein Standard Mix, 15–600 kDa (Sigma-Aldrich) was eluted under the same conditions, and the molecular weights of the protein binder samples were mapped onto the standard curve to determine protein binder molecular weights.

**Analysis of DB2 by nHDX-MS.** Nano-hydrogen/deuterium exchange mass spectrometry (nHDX-MS) experiments were conducted using a modified, fully automated Chronect HDX extended parallel platform (27)(Trajan Scientific and Medical) coupled to an NCS-3500RS Nano system and an Orbitrap Exploris™

480 mass spectrometer (Thermo Fisher Scientific). For DB2 analysis, the protein stock (pET28a/DB2) solution was diluted to 1 µM using PBS (pH 7.4). Samples were labeled at 4 °C in a $D_2O$-based buffer (80% final $D_2O$ content) for 30, 300, 1,000, 3,000, and 10,000 s, using 3 pmol of DB2 per injection. Full deuteration (FD) controls were prepared by incubating the protein at 45 °C for 2 h, followed by equilibration at 4 °C for an additional 3 h prior to quenching and injection (included as a 30,000 s time point). The labeling reactions were quenched at 0 °C with 1 M glycine hydrochloride (glycine-HCl), 4 M urea, and 200 mM tris(2-carboxyethyl)phosphine (TCEP) at a 1:1 volume ratio of sample to quench (final pH 2.5). Samples were digested online for 3 min at 150 µL/min using a dual-protease pepsin and protease XIII column (1:1 ratio, NovaBioassays). Peptides were separated using a 6-min gradient and analyzed via a data-independent acquisition (DIA) workflow, as described by Jain et al. (28). Deuterium incorporation was quantified using HDExaminer™ PRO (version 1.0.0.181, Trajan Scientific and Medical). Samples were analyzed in triplicate. Deuterium content across all time points was measured for 249 peptides spanning the entire protein sequence, yielding 99% coverage and an average redundancy of 32 (**Figure S18, Dataset 1**). HDX-MS data collection, analysis, and reporting followed community recommendations (29). The high sequence coverage and redundancy enabled the extrapolation of D-uptake to the single-residue level, which was calculated using the integrated Keppel & Weis solver (30). To calculate correlations with REMD simulation data (RMSF, Å) at the single-residue level, D-uptake was normalized to the FD control.

### Supplementary Figures

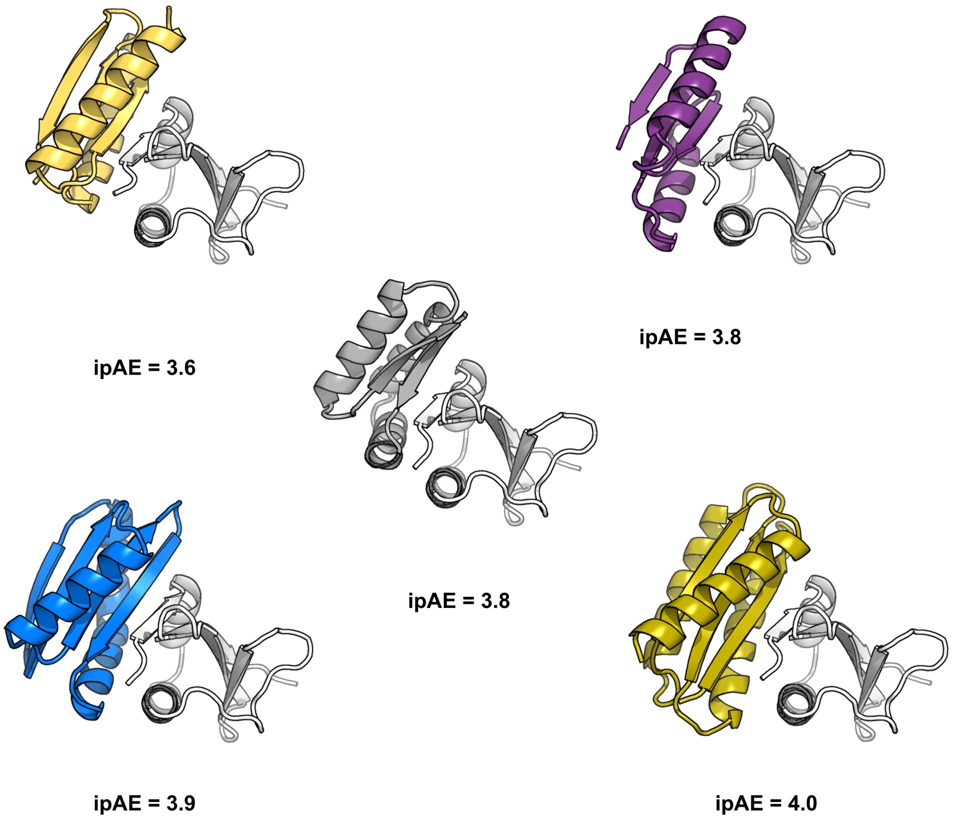


**Figure S1. Structures of the top five designs generated by RFdiffusion without seeds to bind RelE according to AlphaFold2 Initial Guess, with the indicated ipAE.**

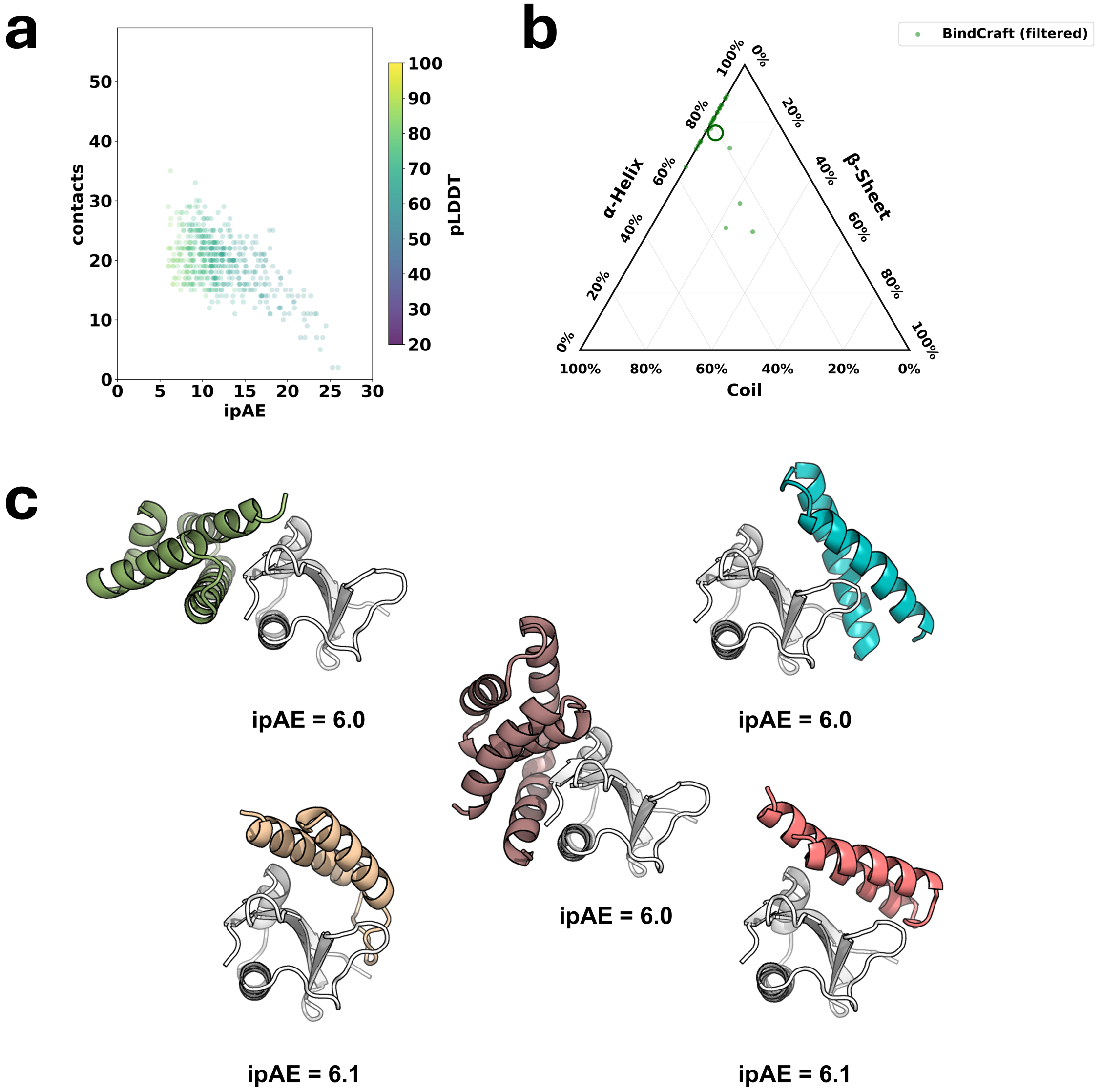


**Figure S2. Structural analysis of designs generated by BindCraft to bind RelE.** (**a**) Scoring metrics after modeling with AF2 IG, showing the relationship between ipAE and contacts (distance < 4 Å) between RelE and the designed protein binder. (**b**) Secondary structure composition of sampled BindCraft designs, calculated by STRIDE, after filtering by ipAE < 10 and contacts ≥ 26. The open circle represents the centroid of the distribution. (**c**) Structures of the top five designs generated by BindCraft using hotspots and default filters according to AF2 IG, with indicated ipAE.

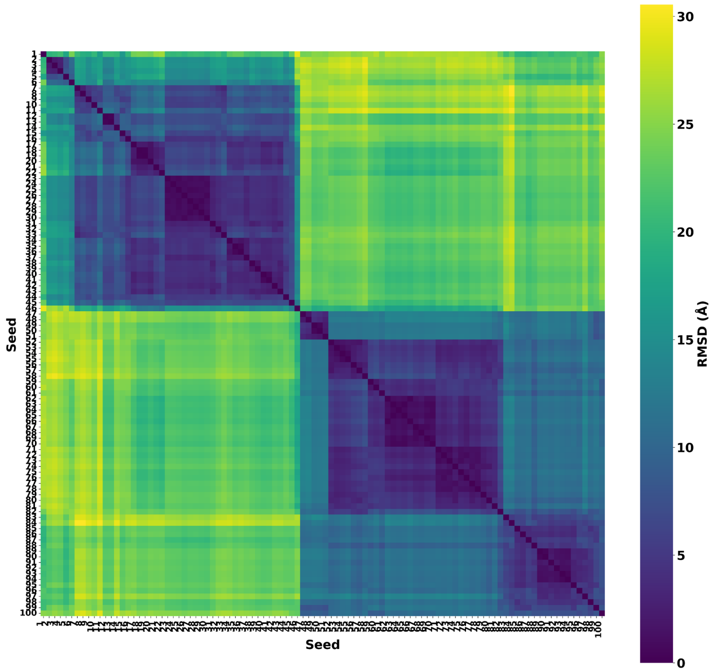


**Figure S3. Inter-seed RMSD analysis of 100 top-scoring seeds generated around RelE.** The RMSD of the backbone atoms of two seeds was calculated after aligning the backbone atoms of RelE. Clustering was performed using a 10 Å RMSD cutoff, resulting in two clusters corresponding to sites 1 and 2, as discussed in the text.

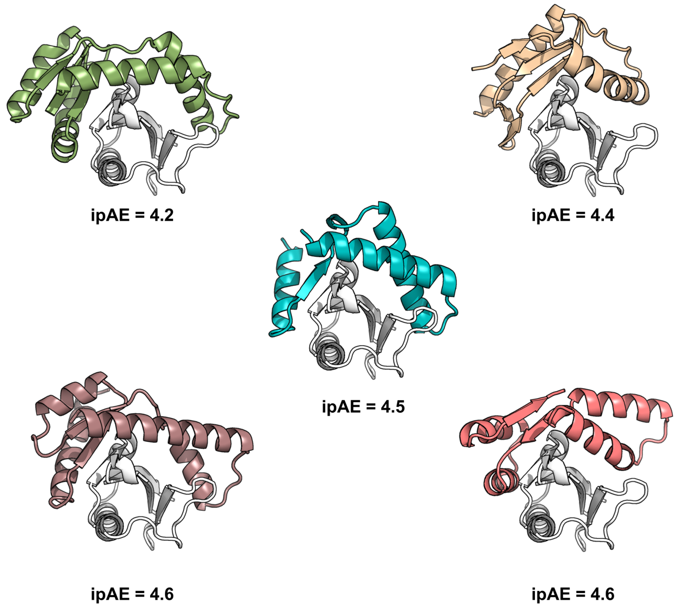


**Figure S4. Structures of the top five designs in the 12,000-design subset generated by seed-guided RFdiffusion, according to AlphaFold2 Initial Guess, with indicated ipAE.**

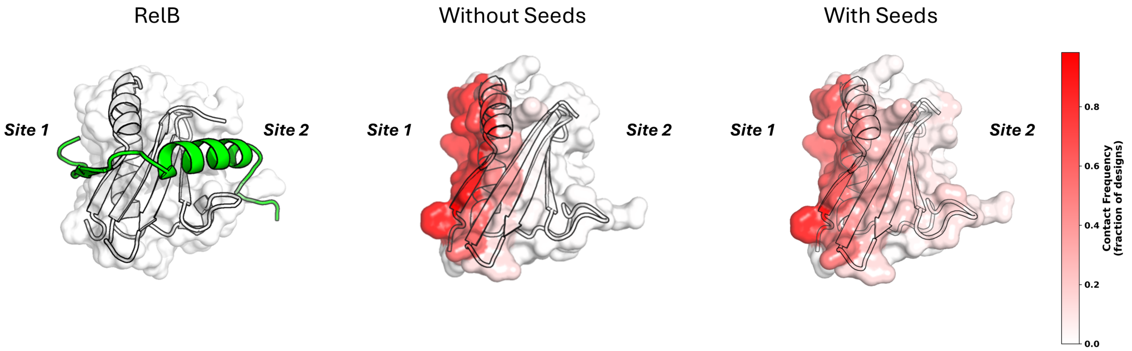


**Figure S5. RelE residue contact frequency among 12,000 designs generated with or without seeds.** The RelB/RelE (PDB: 2KC8) complex is oriented to show both the β-sheet pair and α-helical interaction of RelB with RelE. Using the same orientation, the surface of RelE is colored by the frequency with which 12,000 designs generated without and with seeds contacted each residue in RelE, showing an increased propensity for contacts that wrap around RelE for seed-guided designs.

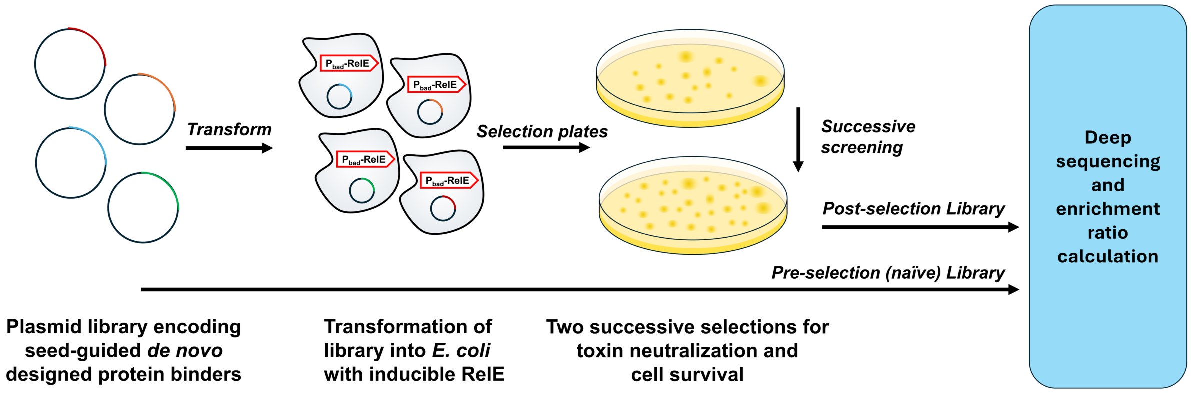


**Figure S6. Survival-based screen for neutralization of RelE toxicity.** Cells expressing RelE under an inducible $P_{bad}$ promoter were transformed with designed RelE binders expressed under the control of $P_{tet}$. Cells that survived were collected, the library was isolated, and the screen was repeated before both post-selection libraries were prepared for deep sequencing. See Supplementary Materials and Methods for details.

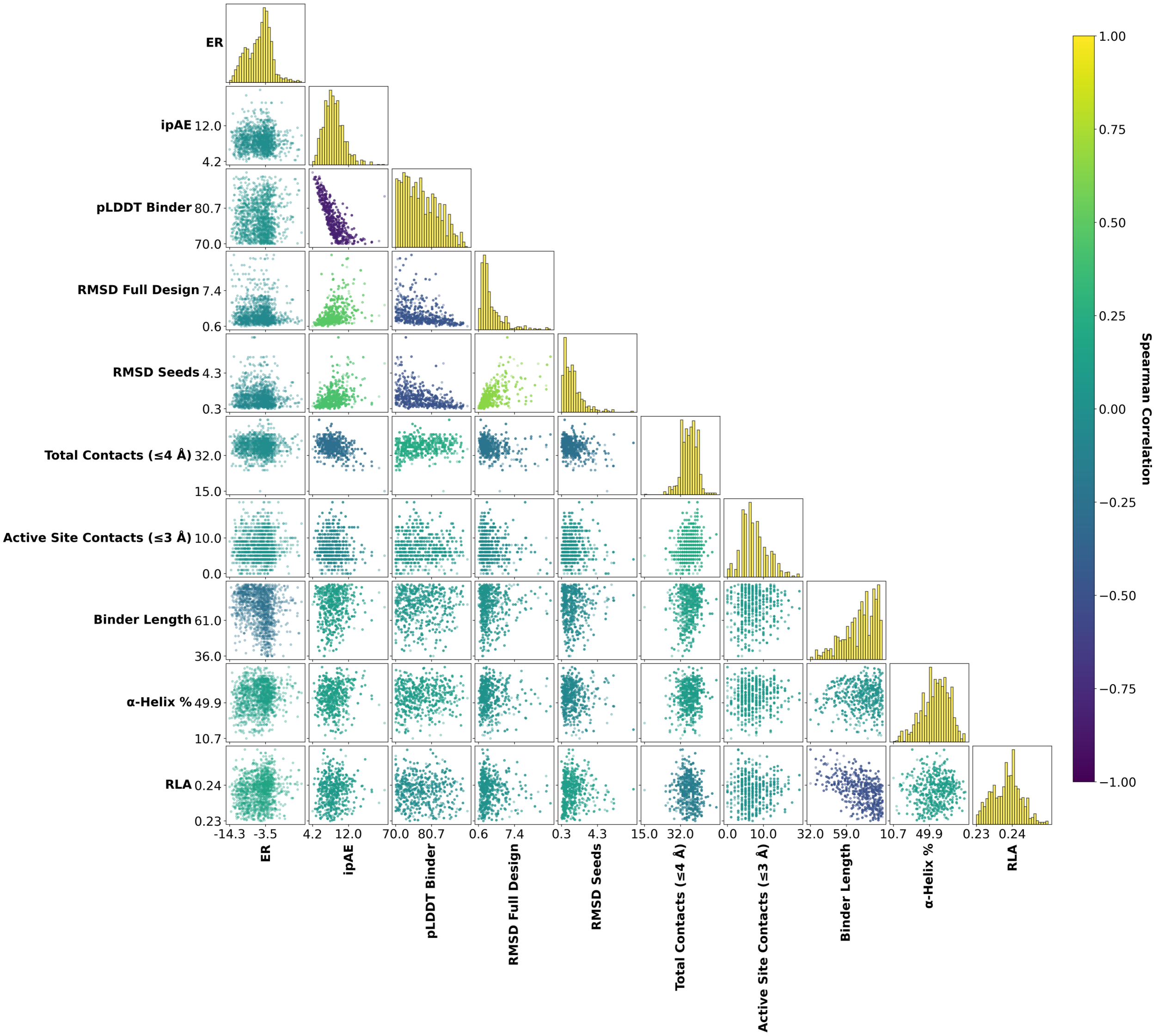


**Figure S7. Correlation of metrics used to score the designs with enrichment ratios in the survival-based screen.** Computational metrics do not correlate strongly with ER values from the second round of screening. The diagonal shows histograms for the metric in the corresponding column.

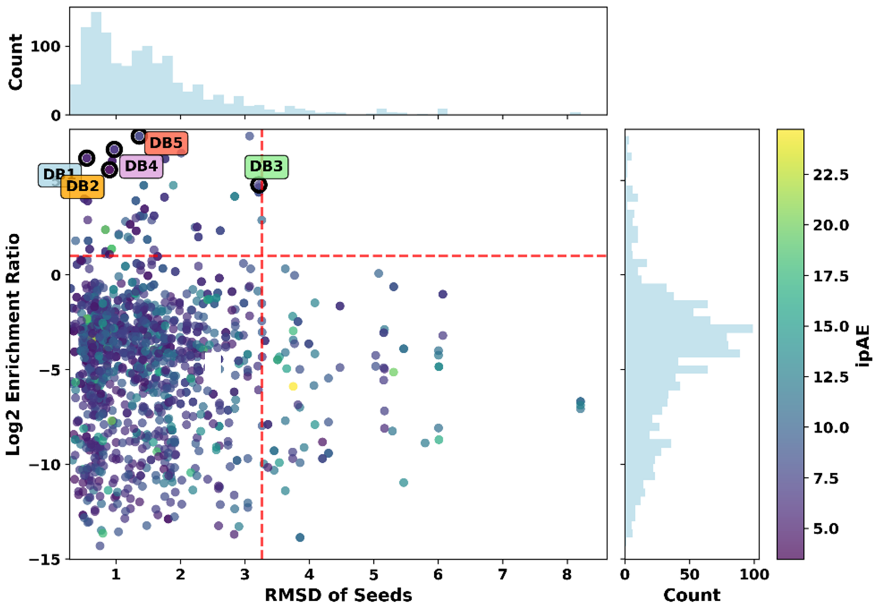


**Figure S8. Comparison of designed binder enrichment ratios (ER) with AlphaFold2 Initial Guess ipAE values and the structural agreement of the modeled designs with the initially placed seeds.** Seed RMSD was calculated between the backbone atoms of seed regions in the RFdiffusion design and the corresponding atoms in the AlphaFold2 Initial Guess structure after aligning RelE. Designs DB1–DB5 are labeled.

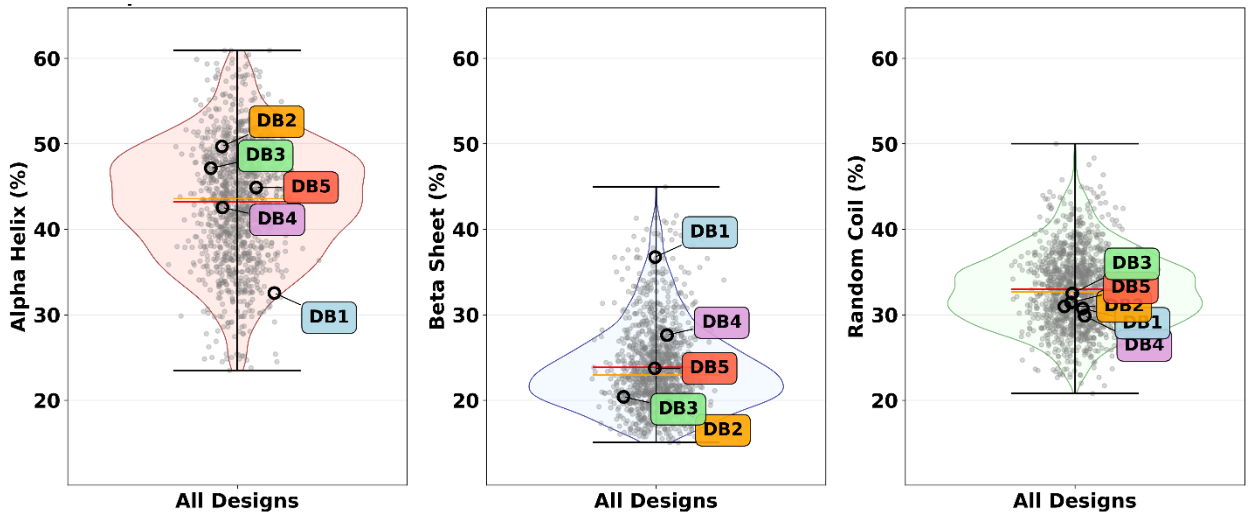


**Figure S9. Secondary structure analysis for the designed binder library.** Secondary structure was calculated by STRIDE for designed binders that were tested in experimental screening, highlighting DB1–DB5.

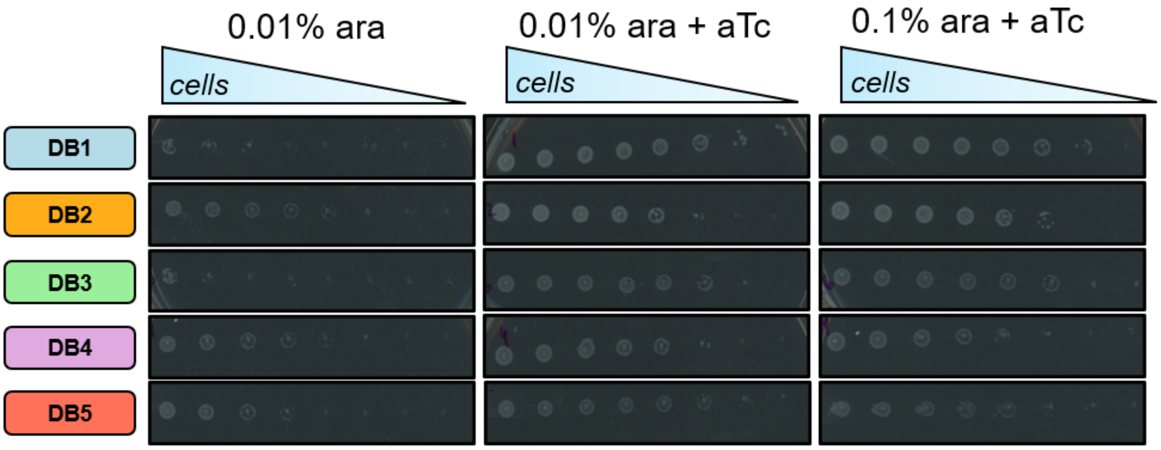


**Figure S10. DB1–DB5 activity assessed by toxicity spotting assay.** Expression of DB1–DB5, induced by aTc, rescued growth when RelE expression was induced by 0.01% or 0.1% arabinose.

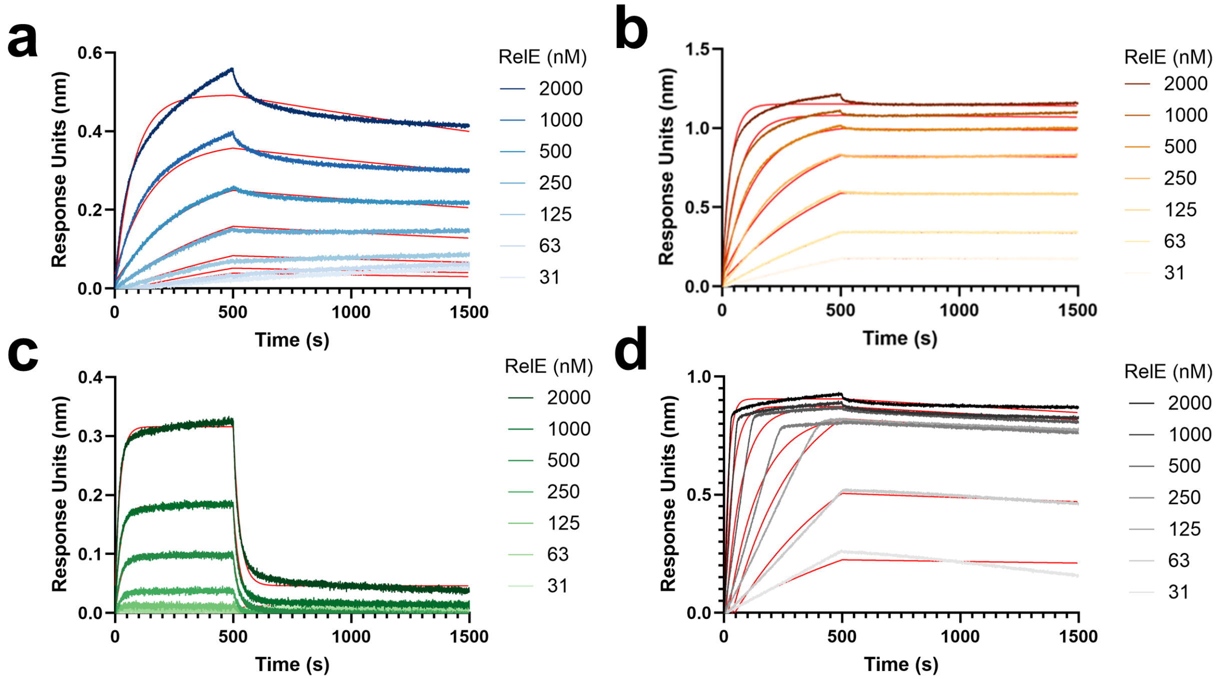


**Figure S10. Representative biolayer interferometry (BLI) traces for RelE binding to designed proteins**. Traces and fits of RelE binding to surface-immobilized (**a**) DB1, (**b**) DB2, (**c**) DB3, and (**d**) $RelB_{pep}$, showing fits used to determine the values in **Table 1**.

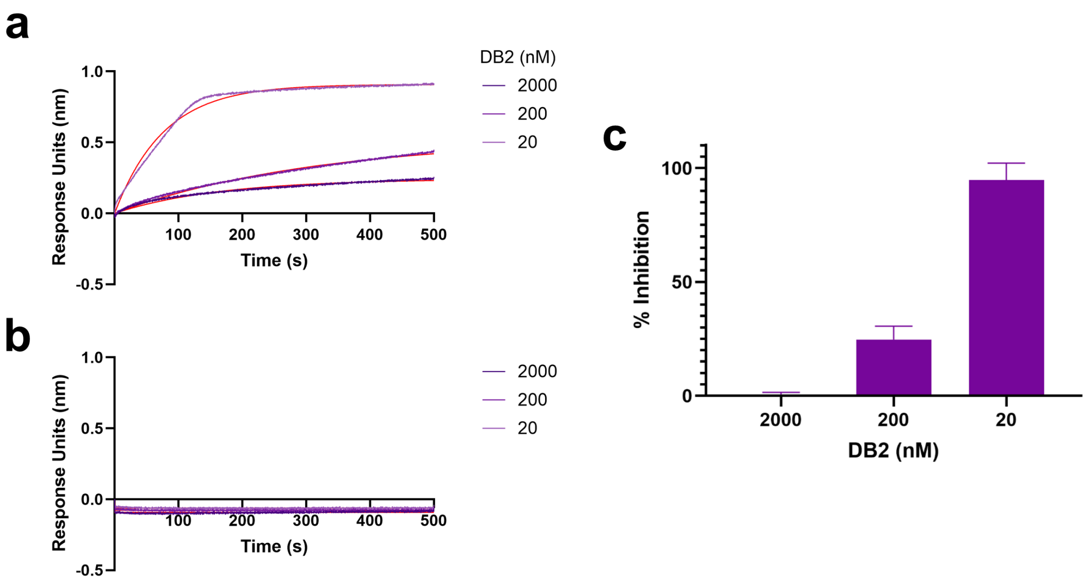


**Figure S11. Competitive binding of RelB$_{pep}$ and DB2 assessed by BLI.** Representative biolayer interferometry traces (purple) demonstrating binding of RelE, but not DB2, to RelBpep. RelB$_{pep}$ was immobilized and incubated with (**a**) 500 nM RelE plus DB2 at concentrations of 20–2,000 nM or (**b**) DB2 at concentrations of 20–2,000 nM. (**c**) Inhibition was calculated using the average reduction in the fitted maximum signal (from kinetic fits, shown in red) relative to RelE binding in the absence of DB2. Error bars represent the standard deviation of two independent trials for both bound and unbound assays from separate DB2 and RelB$_{pep}$ preparations.

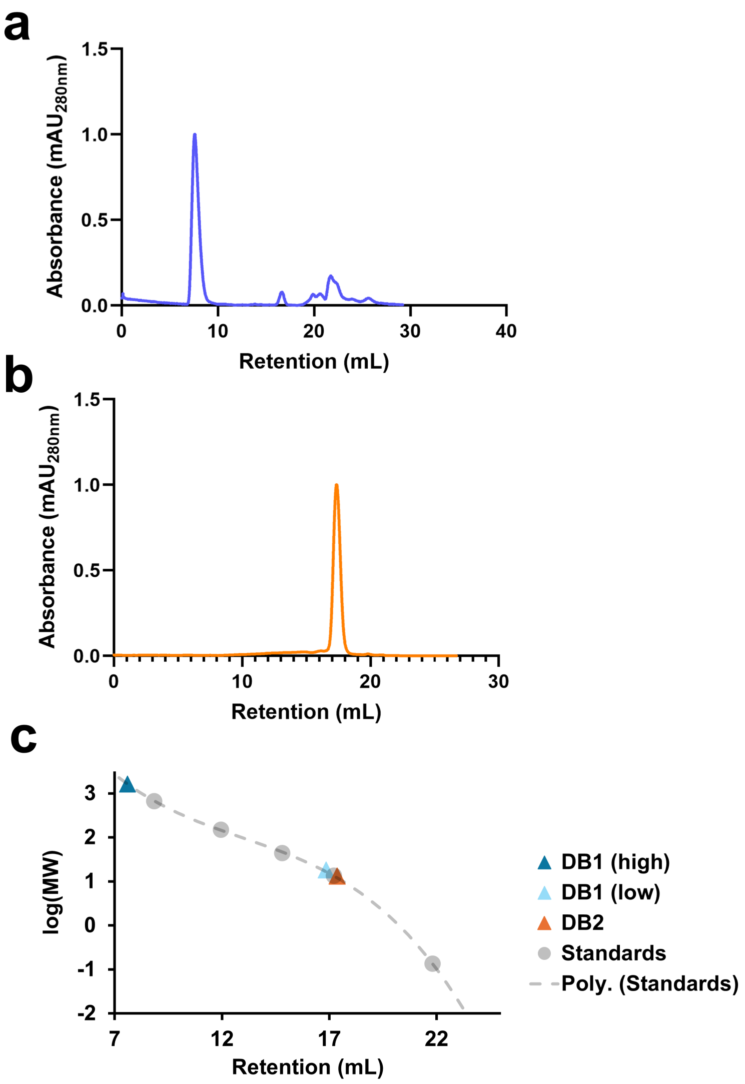


**Figure S12. DB1 aggregates, whereas DB2 elutes as a monomer in size-exclusion chromatography. Elution profiles of (a) DB1 and (b) DB2.** (c) Logarithmic mapping of retention time using Cytiva standards on a Superdex 200 GL column. DB1 and DB2 retention volumes are shown relative to standards. DB2 (14 kDa) exhibited a monodisperse peak corresponding to ~13 kDa. DB1 (15 kDa) exhibited a peak at a MW of ~1.6 MDa and a small peak at ~18 kDa. The large peak is consistent with the poor solubility of DB1 and its propensity to aggregate.

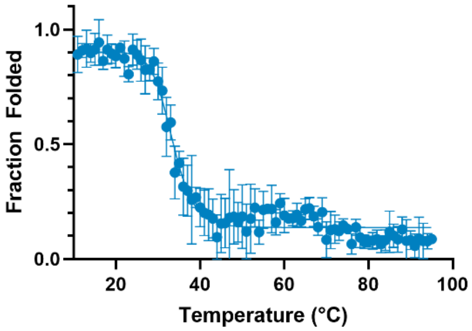


**Figure S13. Thermal stability of DB1.** Fraction of folded DB1, based on CD signal at 222 nm, from 10 to 95 °C. Error bars represent the standard deviation of three replicate experiments.

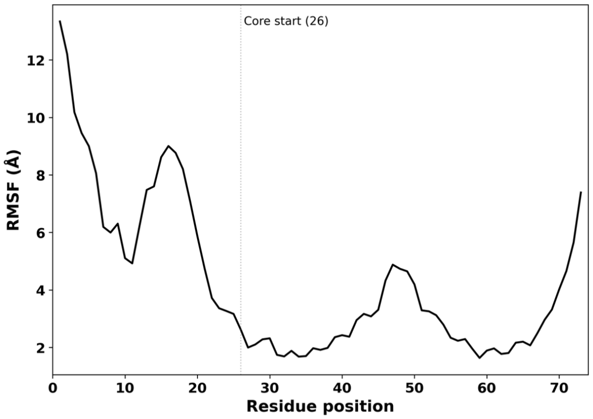


**Figure S14. Per-residue flexibility of DB2 in REMD simulations initiated from the bound conformation.** Per-residue Cα root mean square fluctuation (RMSF) of the 0–500 ns demultiplexed 300 K trajectory. The N-terminal region (residues 1–25) exhibited high flexibility (RMSF up to ~13 Å), while the structured core (residues 26–73, demarcated by a vertical line) maintained low fluctuations.

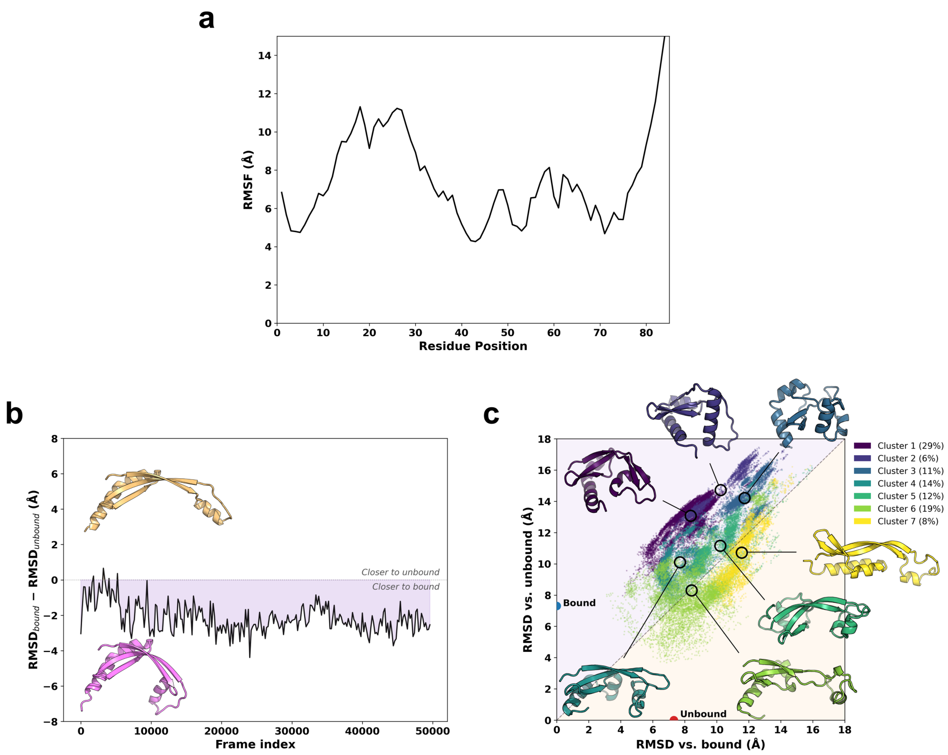


**Figure S15. DB1 exhibits large conformational changes over the course of a 500-ns REMD simulation. (a)** Per-residue Cα root mean square fluctuations (RMSF) of the 0–500 ns demultiplexed 300 K trajectory of DB1. (**b**) REMD simulation of DB1 showed modestly greater similarity to the predicted bound conformation (purple) compared with the unbound conformation (orange). $RMSD_{bound} - RMSD_{unbound}$ values, averaged over 200-frame windows for a simulation temperature of 300 K, are plotted as a function of simulation time. (**c**) Structures from the REMD simulation were projected based on their RMSD relative to the bound and unbound AlphaFold3 reference structures. Seven clusters identified by k-means clustering on backbone Cα coordinates are colored by a gradient from most bound-like (Cluster 1, dark) to most unbound-like (Cluster 7, bright). Representative structures corresponding to the centroid for each cluster are shown. Red and blue dots indicate the positions of the unbound and bound AlphaFold3 reference structures, respectively. Orange and purple shading denote greater similarity to the predicted unbound and bound structures, respectively.

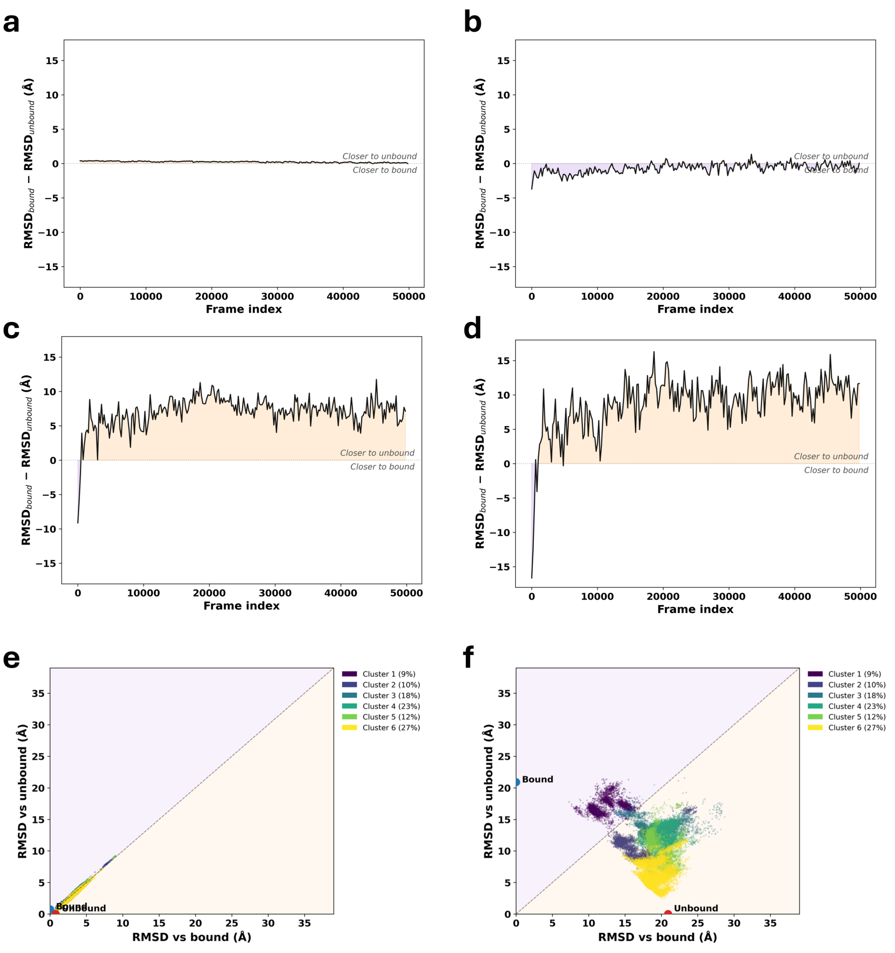


**Figure S16. Progression of the REMD simulation of DB2 initiated from the bound-state conformation.** (**a**–**b**) $RMSD_{bound}$ − $RMSD_{unbound}$ values, averaged over 200-frame windows for a simulation temperature of 300 K, are plotted as a function of simulation time. (**a**) Analysis of the C-terminal core (residues 26–73) after aligning on the C-terminal core and (**b**) the N-terminal region (residues 1–25) after aligning on the N-terminal region, with orange and purple shading indicating conformations more similar to the unbound and bound states, respectively. Both regions individually maintain low RMSD from the AlphaFold3-predicted structures, though the N-terminal region exhibits greater structural fluctuations. (**c**) The N-terminal region (residues 1–25) after aligning on the C-terminal core and (**d**) the C-terminal core (residues 26–73) after aligning on the N-terminal region. In both cross-region analyses, the non-aligned region shows large displacements, indicating the two regions undergo significant relative motion while each individually retains its local fold. (**e**–**f**) Two-dimensional RMSD landscape of the ensembles projected onto backbone RMSD relative to the bound and unbound reference structures using alignment on the C-terminal core. (**e**) RMSD landscape of the C-terminal helices. (**f**) RMSD landscape of the N-terminal helix. Clusters correspond to clustering using the full-backbone RMSD landscape shown in **Figure 5**. Red and blue dots indicate the positions of the unbound and bound AlphaFold3 reference structures, respectively. Orange and purple shading denote regions more similar to the unbound and bound states, respectively.

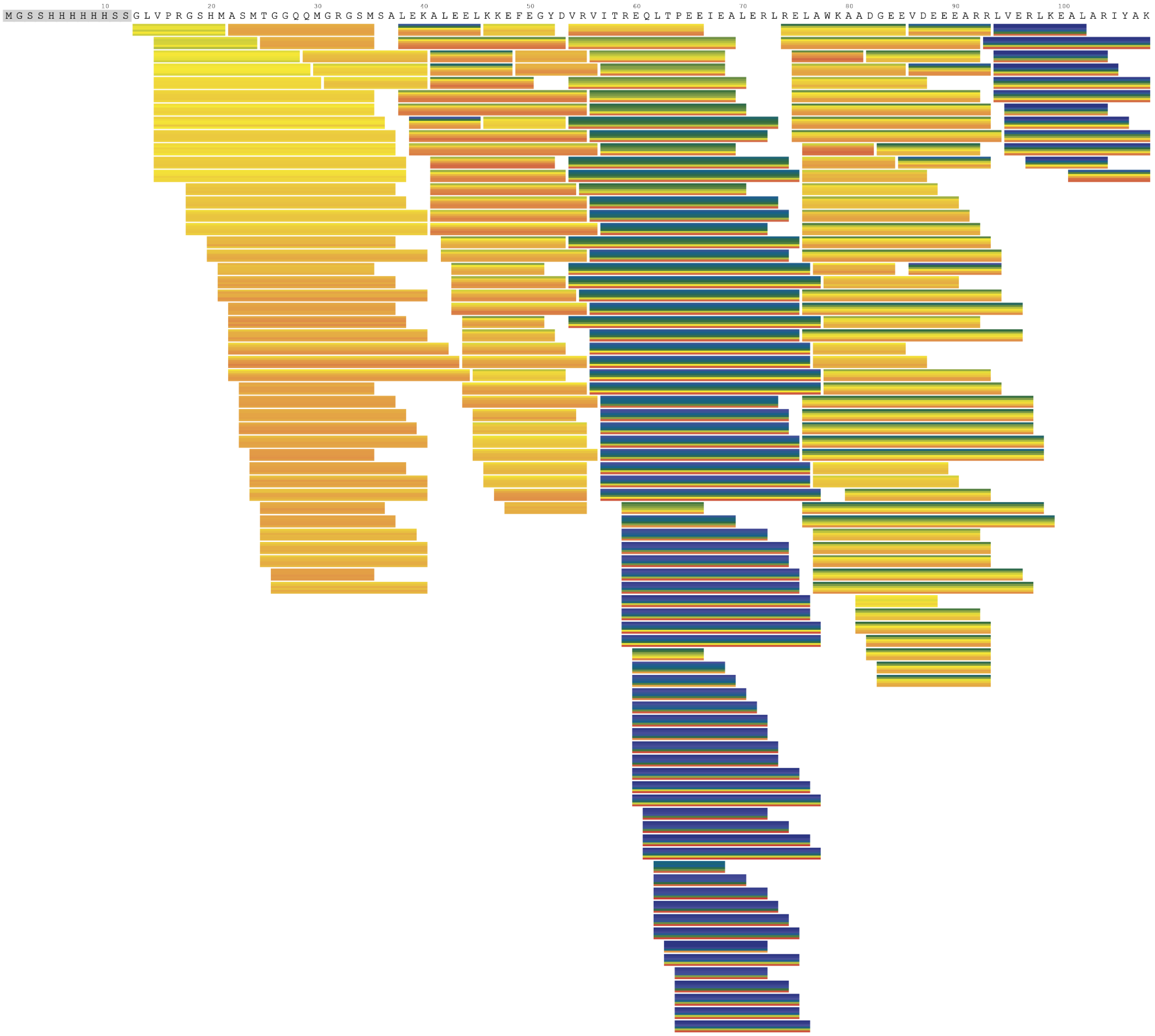


**Figure S17. Deuterium uptake of DB2 at the peptide level.** D-uptake across all time points was measured for 249 peptides, resulting in 99% coverage and an average redundancy of 32. His-tag peptides were removed due to rapid back-exchange. Data are not normalized to the fully deuterated control (30,000 s time point).

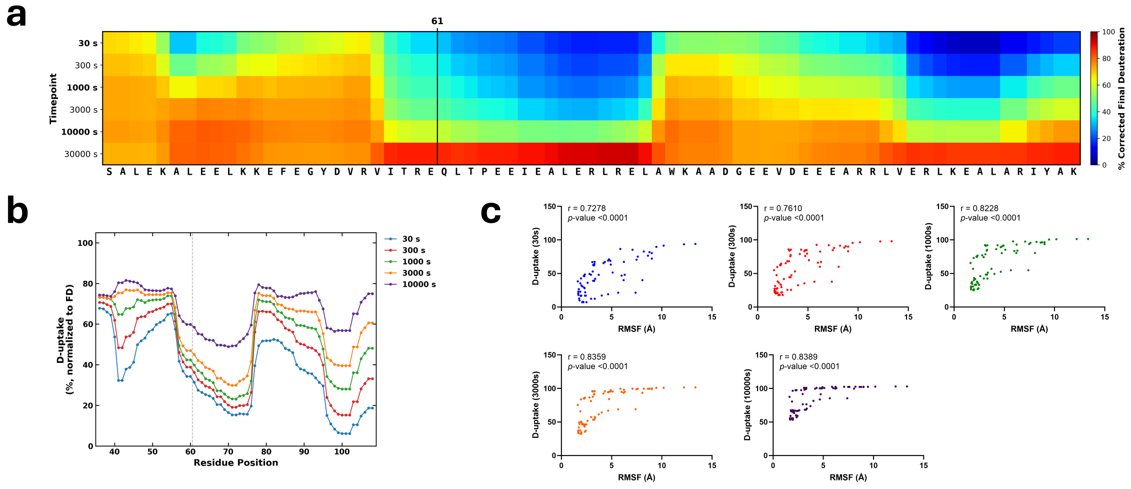


**Figure S18. Single-residue deuterium uptake kinetics of DB2.** (**a**) Single-residue-resolved HD exchange heatmap of DB2 across multiple time points calculated using the Keppel & Weis solver (30). His-tag peptides were removed from the final analysis due to rapid exchange. (**b**) The data from panel **a** are replotted with normalization to a fully deuterated control (30,000 s). (**c**) Spearman correlation analysis between FD-normalized single-residue D-uptake and RMSF values (Å) from REMD simulations for five time points. The correlation analysis used the entire protein binder (DB2 residues 1–73).

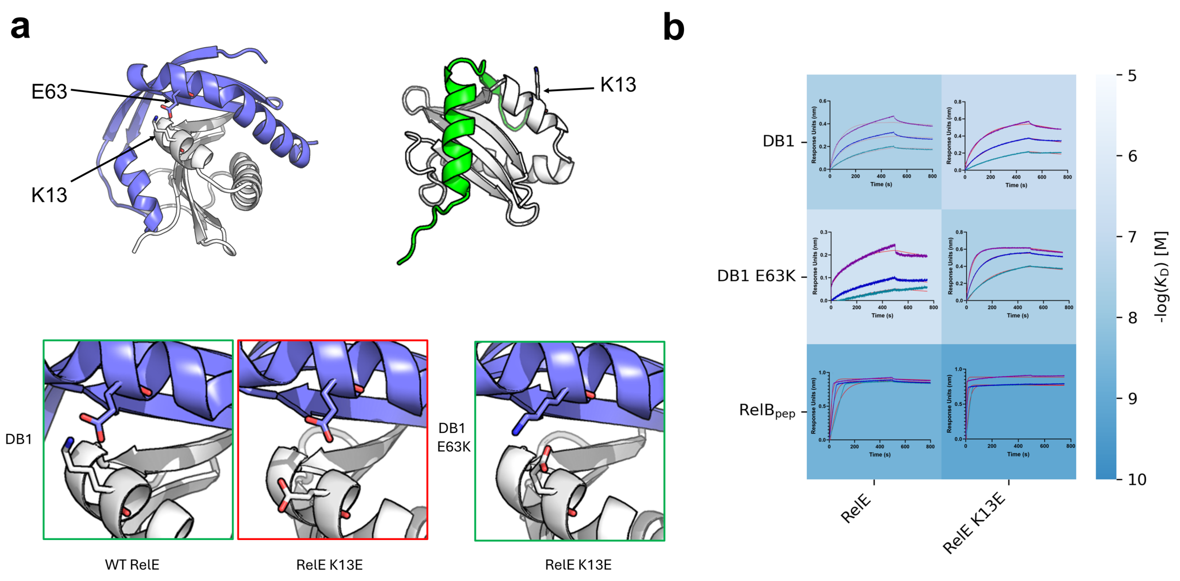


**Figure S19. Mutational analysis of the predicted interface between DB1 and RelE. (a)** AlphaFold3-predicted structure of DB1 bound to RelE and the crystal structure of $RelB_{pep}$ bound to RelE (PDB: 2KC8), indicating sites of charge-swap mutations. Boxes show modeled structures for mutants with oppositely charged residue pairs (green) or like-charged residue pairs (red). (**b**) Heatmap of average $-\log(K_D)$ measured by BLI (M), with red and green boxes highlighting charge pairs as in panel **a**. Binding affinity is reported as the average of two independent replicate experiments. Representative binding curves and fits are shown within the corresponding matrix cells. Fitted kinetic parameters are reported in Table 1.

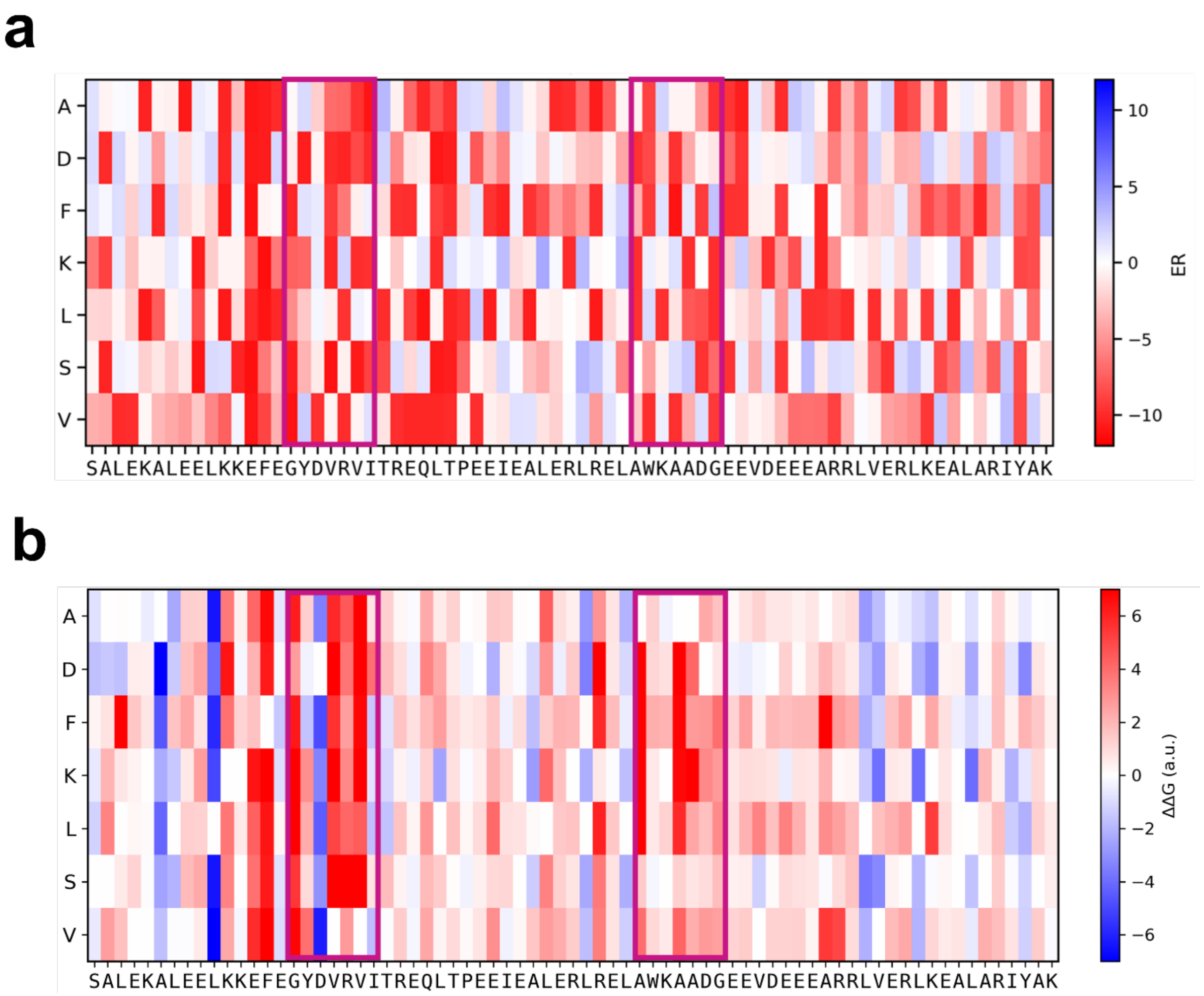


**Figure S20. Effects of mutations on DB2 activity and predicted binding to RelE.** Each cell shows data for a point mutant of DB2. Positions are indicated by columns, with the sequence of DB2 included at the bottom. Rows indicate the residue that was substituted. (**a**) ER values from the growth-based screen. (**b**) ΔΔGbind values computed with PottsMPNN using the AlphaFold3-predicted structure of DB2.

### Supplementary Tables

#### Table S1. Sequences of designed binders, RelB$_{pep}$, and RelE.

| Construct | Vector | MW (kDa) | Sequence |
|---|---|---|---|
| DB1 [a] | pKVS45 | 10.9 | MPTVTKSTETRTFKGVTLEEMYKALEKDGIESDKTETVTKTVLTVEIAPGVYLRISASEADLEEAEAAL PELYAELKAELEAKKK |
| DB2 [a] | pKVS45 | 10.0 | MSALEKALEELKKEFEGYDVRVITREQLTPEEIEALERLRELAWKAADGEEVDEEEARRLVERLKEAL ARIYAK |
| DB3 [a] | pKVS45 | 9.7 | MGAAAAAAALAAKVEEARERLRSLINIDDFSTWTEEEKKELAELLELYLANAKVVRVDLSKGLEEGIEK AIEEAEK |
| DB4 [a] | pKVS45 | 7.2 | MSIGPIEIDGLKVDRVVPTTPEVEELRKKAEEALERTDEEEERRRRLEELLKE |
| DB5 [a] | pKVS45 | 10.3 | MMEQKVVFSEGVSEEDKELAKELGKELADKAPEVRVLTREEAQKLLKGVSLEELLRGIEEEERRERE ERRKRREE |
| DB1 | pDW363 | 28.6 | MAGGLNDIFEAQKIEWHEDTGGSSHHHHHHGSGSGSDSEVNQEAKPEVKPEVKPETHINLKVSDGS SEIFFKIKKTTPLRRLMEAFAKRQGKEMDSLTYFLYDGIEIQADQTPEDLDMEDNDIIEAHREQIGGSG SGMPTVTKSTETRTFKGVTLEEMYKALEKDGIESDKTETVTKTVLTVEIAPGVYLRISASEADLEEAEA ALPELYAELKAELEAKKK |
| DB2 | pDW363 | 27.7 | MAGGLNDIFEAQKIEWHEDTGGSSHHHHHHGSGSGSDSEVNQEAKPEVKPEVKPETHINLKVSDGS SEIFFKIKKTTPLRRLMEAFAKRQGKEMDSLTYFLYDGIEIQADQTPEDLDMEDNDIIEAHREQIGGSG SGMSALEKALEELKKEFEGYDVRVITREQLTPEEIEALERLRELAWKAADGEEVDEEEARRLVERLKE ALARIYAK |
| DB3 | pDW363 | 27.4 | MAGGLNDIFEAQKIEWHEDTGGSSHHHHHHGSGSGSDSEVNQEAKPEVKPEVKPETHINLKVSDGS SEIFFKIKKTTPLRRLMEAFAKRQGKEMDSLTYFLYDGIEIQADQTPEDLDMEDNDIIEAHREQIGGSG SGMGAAAAAAALAAKVEEARERLRSLINIDDFSTWTEEEKKELAELLELYLANAKVVRVDLSKGLEEGI EKAIEEAEK |
| DB1 E63K | pDW363 | 28.6 | MAGGLNDIFEAQKIEWHEDTGGSSHHHHHHGSGSGSDSEVNQEAKPEVKPEVKPETHINLKVSDGS SEIFFKIKKTTPLRRLMEAFAKRQGKEMDSLTYFLYDGIEIQADQTPEDLDMEDNDIIEAHREQIGGSG SGMPTVTKSTETRTFKGVTLEEMYKALEKDGIESDKTETVTKTVLTVEIAPGVYLRISASEADLEKAEA ALPELYAELKAELEAKKK |
| DB2 R20E | pDW363 | 27.7 | MAGGLNDIFEAQKIEWHEDTGGSSHHHHHHGSGSGSDSEVNQEAKPEVKPEVKPETHINLKVSDGS SEIFFKIKKTTPLRRLMEAFAKRQGKEMDSLTYFLYDGIEIQADQTPEDLDMEDNDIIEAHREQIGGSG SGMSALEKALEELKKEFEGYDVEVITREQLTPEEIEALERLRELAWKAADGEEVDEEEARRLVERLKE ALARIYAK |
| RelB$_{pep}$ | pDW363 | 22.3 | MAGGLNDIFEAQKIEWHEDTGGSSHHHHHHGSGSGSDSEVNQEAKPEVKPEVKPETHINLKVSDGS SEIFFKIKKTTPLRRLMEAFAKRQGKEMDSLTYFLYDGIEIQADQTPEDLDMEDNDIIEAHREQIGGSG SGMKQTLLSDEDAELVEIVKERLRNPKPVRVTLDEL |
| DB1 | pET28a | 15.0 | MGSSHHHHHHSSGLVPRGSHMASMTGGQQMGRGSMRGSHHHHHHENLYFQMPTVTKSTETRTF KGVTLEEMYKALEKDGIESDKTETVTKTVLTVEIAPGVYLRISASEADLEEAEAALPELYAELKAELEAK KK |
| DB2 | pET28a | 14.2 | MGSSHHHHHHSSGLVPRGSHMASMTGGQQMGRGSMSALEKALEELKKEFEGYDVRVITREQLTPE EIEALERLRELAWKAADGEEVDEEEARRLVERLKEALARIYAK |
| RelE | pET28a | 19.2 | MGSSHHHHHHSSGLVPRGSHMASMTGGQQMGRGSMRGSHHHHHHENLYFQMAYFLDFDERALK EWRKLGSTVREQLKKKLVEVLESPRIEANKLRGMPDCYKIKLRSSGYRLVYQVIDEKVVVFVISVGKA EASEVYSEAVKRIL |
| RelE D6K | pET28a | 14.8 | MRGSHHHHHHHHGSGMAYFLKFDERALKEWRKLGSTVREQLKKKLVEVLESPRIEANKLRGMPDC YKIKLRSSGYRLVYQVIDEKVVVFVISVGKAEASEVYSEAVKRIL |
| RelE K13E | pET28a | 14.8 | MRGSHHHHHHHHGSGMAYFLDFDERALEEWRKLGSTVREQLKKKLVEVLESPRIEANKLRGMPDC YKIKLRSSGYRLVYQVIDEKVVVFVISVGKAEASEVYSEAVKRIL |

[a] Sequence matches the designed sequence from the computational pipeline with an additional N-terminal methionine from the start codon; this residue is excluded from the residue numbering used in the computational analyses.

**Table S2. Properties of designed binders.**

| Binder | Enrichment Ratio[a] | ipAE[b] | pLDDT[c] | Seed RMSD[d] (Å) | Helicity[e] (%) |
|---|---|---|---|---|---|
| DB1 | 6.2 | 5.3 | 88 | 0.6 | 32.5 |
| DB2 | 5.5 | 3.5 | 94 | 0.9 | 49.7 |
| DB3 | 4.8 | 7.1 | 81 | 3.2 | 47.1 |
| DB4 | 6.6 | 6.3 | 87 | 1.0 | 42.5 |
| DB5 | 7.3 | 7.2 | 83 | 1.4 | 44.9 |

[a] Sequence enrichment ratio of the binder in the post-selection pool relative to the naïve library from bacterial screening against RelE.

[b] Interface predicted aligned error (Å) from the AlphaFold2 Initial Guess prediction of the binder–RelE complex; lower values indicate higher confidence in the predicted interface geometry.

[c] Predicted local distance difference test (0–100) from the AlphaFold2 Initial Guess prediction of the binder–RelE complex, averaged over the binder chain.

[d] Backbone RMSD (Å) between the seed regions in the RFdiffusion design and the corresponding residues in the AlphaFold2 Initial Guess-evaluated complex.

[e] Predicted helicity of the AlphaFold3-predicted binder structure (fraction of residues assigned to α-helix by STRIDE).

**Table S3. Circular dichroism spectroscopy analysis of DB1 and DB2.**

| | | *Circular dichroism (BeStSel deconvolution)* | | |
|---|---|---|---|---|
| **Binder** | **T (°C)** | **α-helix**[a] (%) | **β-sheet**[a] (%) | **Random coil**[a] (%) |
| DB1[b] | 10 | 1 ± 2 (1 ± 3) | 52 ± 1 (74 ± 1) | 47 ± 2 |
| | 25 | 1 ± 2 (1 ± 3) | 52 ± 3 (74 ± 4) | 47 ± 1 |
| | 45 | 0 ± 0 (0 ± 0) | 55 ± 3 (78 ± 4) | 45 ± 3 |
| | 65 | 1 ± 1 (1 ± 1) | 48 ± 3 (68 ± 4) | 52 ± 3 |
| | 85 | 2 ± 4 (3 ± 6) | 44 ± 4 (62 ± 6) | 54 ± 2 |
| DB2[b] | 10 | 54 ± 4 (80 ± 6) | 12 ± 4 (18 ± 6) | 34 ± 1 |
| | 25 | 51 ± 3 (75 ± 4) | 13 ± 2 (19 ± 3) | 36 ± 2 |
| | 45 | 47 ± 4 (70 ± 6) | 15 ± 3 (22 ± 4) | 38 ± 3 |
| | 65 | 42 ± 3 (62 ± 4) | 16 ± 4 (24 ± 6) | 42 ± 1 |
| | 85 | 41 ± 1 (61 ± 1) | 16 ± 2 (24 ± 3) | 44 ± 1 |

| | | *Replica-exchange molecular dynamics*[c] *helicity of DB2* | | |
|---|---|---|---|---|
| **Cluster** | **Frames (*n*)** | **Full** (% α-helix) | **N-term (1–25)** (% α-helix) | **Core (26–73)** (% α-helix) |
| 1 | 855 | 57.9 ± 6.2 | 43.7 ± 7.3 | 66.4 ± 8.4 |
| 2 | 902 | 56.9 ± 4.5 | 44.9 ± 8.1 | 64.3 ± 6.7 |
| 3 | 1,634 | 61.2 ± 4.4 | 39.8 ± 11.7 | 73.6 ± 4.3 |
| 4 | 2,428 | 62.5 ± 5.2 | 45.4 ± 9.1 | 72.6 ± 6.0 |
| 5 | 1,130 | 60.1 ± 7.0 | 36.8 ± 19.6 | 73.5 ± 5.9 |
| 6 | 2,051 | 62.6 ± 5.4 | 43.9 ± 12.9 | 73.7 ± 4.3 |
| **Overall** | **10,000** | **61.2 ± 5.7** | **42.8 ± 12.1** | **72.1 ± 6.5** |

[a] Mean ± s.d. of three replicate wavelength scans at each temperature, as calculated by BeStSel (DB2 at 85 °C: n = 2).

[b] Values in parentheses are corrected to a design-only basis, assuming structural signal arises only from the design sequence and not the construct tag: corrected % = raw % × (full-construct length / design length). DB1: 119/84; DB2: 108/73.

[c] A 500-ns replica-exchange molecular dynamics trajectory of the DB2 design sequence (no construct tags). The 10,000 frames were partitioned into six clusters; per-residue α-helix content was calculated by STRIDE and averaged within each cluster, with the Overall row reporting the population-weighted ensemble mean. N-term: residues 1–25. Core: residues 26–73.
